# Subcircuits of deep and superficial CA1 place cells support efficient spatial coding across heterogeneous environments

**DOI:** 10.1101/2020.04.17.047399

**Authors:** Farnaz Sharif, Behnam Tayebi, György Buzsáki, Sebastien Royer, Antonio Fernandez-Ruiz

**Author notes:** Correspondence: Antonio Fernandez-Ruiz or Sebastien Royer.

## Abstract

The hippocampus is thought to guide navigation by forming a cognitive map of space. However, the behavioral demands for such a map can vary depending on particular features of a given environment. For example, an environment rich in cues may require a finer resolution map than an open space. It is unclear how the hippocampal cognitive map adjusts to meet these distinct behavioral demands. To address this issue, we examined the spatial coding characteristics of hippocampal neurons in mice and rats navigating different environments. We found that CA1 place cells located in the superficial sublayer were more active in cue-poor environments, and preferentially used a firing rate code driven by intra-hippocampal inputs. In contrast, place cells located in the deep sublayer were more active in cue-rich environments and expressed a phase code driven by entorhinal inputs. Switching between these two spatial coding modes was supported by the interaction between excitatory gamma inputs and local inhibition.

## INTRODUCTION

Animals navigate different environments in search of food, shelter, or to escape threats. To successfully accomplish these tasks, they must be able to orient themselves based on environmental landmarks or using path-integration. The hippocampal formation is believed to support this process by creating a ‘cognitive map’ of the animal’s travel within its environment (O’Keefe and Nadel, 1978). Hippocampal pyramidal cells that fire selectively when the animal is at a particular location, known as ‘place cells’ (O’Keefe and Dostrovsky, 1971), are considered the building blocks of this cognitive map. Different place cells fire at distinct locations (i.e., in their ‘place fields’) such that the whole environment is represented by the combined activity of the population of place cells (O’Keefe, 1976). The correlation between the firing rate of a place cell and position is referred as the ‘rate code’, since it is possible for an external observer (and also for a downstream ‘reader’ structure) to accurately decode the animal’s position from the firing rates of the place cell population (Wilson and McNaughton, 1993).

The firing patterns of hippocampal place cell are globally coordinated by theta oscillations, which synchronize place cells with overlapping place fields into functional ‘assemblies’ and represent the animal’s trajectory by linking assembly sequences that encode previous, current and future positions (i.e., in ‘theta sequences’; Skaggs et al., 1996; Dragoi and Buzsaki, 2006; Foster and Wilson, 2007; Gupta et al., 2012). The spike timing of individual place cells advances with respect to the ongoing theta rhythm as an animal moves through the neuron’s place field (‘phase precession’; O’Keefe and Recce 1993), and thus position can also be decoded from theta phase (a ‘phase code’). Yet, it is not clear how these complementary codes for space are used interchangeably during navigation (Harris et al., 2002; Mehta et al., 2002; Huxter et al., 2003; Huxter et al., 2008; O’Keefe and Burgess, 2005; Cei et al., 2014; Lasztoczi and Klausberger, 2016; Fernandez-Ruiz et al., 2017).

An advantage of the phase code is that it has higher spatial resolution than the rate code (Jensen and Lisman, 2000; Huxter et al., 2008; Tingley and Buzsaki, 2018). On a coarse spatial scale (i.e. tens of cm to meters), larger than the average place field size, a rate code is reliable since place cell firing outside its fields is minimal. However, since the firing rate of a place cell increases and decreases as the animal crosses the field, identical firing rates can correspond to positions separated up to tens of cm. On the other hand, theta-phase of place cell spikes shows a unidirectional advancement along the place field, offering a more reliable mechanism to encode position at a finer spatial scale.

A limitation shared by most previous studies is that they analyzed hippocampal spatial coding while animals were exploring cue-impoverished environments such as linear tracks or open fields, in which the animal’s behavior did not require discrimination at finer spatial scales. Studies that assessed the modulation of hippocampal place cells by sensory cues, objects and landmarks, reported heterogeneous responses (Knierim et al., 1998; Save et al., 2000; Knierim et al., 2002; Burke et al., 2011). This heterogeneity of responses has been attributed to the anatomical location of the place cells recorded (Royer et al., 2010; Igarashi et al., 2014; Danielson et al., 2016; Geiller et al., 2017). In the CA1 region in particular, two populations of pyramidal cells have been distinguished based on their radial anatomical position (Lorente de No, 1934; Slomianka et al., 2011). CA1 pyramidal cells located closer to the stratum oriens (or ‘deep’ sublayer) and those located closer to the stratum radiatum (‘superficial’ sublayer) have striking genetic, anatomical and physiological differences (Mizuseki et al., 2011; Lee et al., 2014; Maroso et al., 2016; Valero et al., 2015; Oliva et al., 2016a; Cembroski et al., 2016). We hypothesize that the heterogeneity of CA1 pyramidal cells confers the hippocampus a larger computational flexibility that could support spatial coding across different environments.

Based on this background, we hypothesized that in situations in which a coarser spatial map is sufficient to support behavioral demands, such as a cue-impoverished environment, the rate code dominates. In contrast, in feature-rich environments, where a finer resolution map is required, phase code is preferred. To test this hypothesis, we recorded hippocampal place cells in mice and rats navigating cue-poor and cue-rich environments and found that superficial and deep CA1 place cells were mainly recruited in cue-poor and local cue-rich mazes, respectively. In cue-poor environments, place cells were more spatially selective and their firing rates carried more spatial information. In contrast, in the presence of local cues (objects) the spatial rate code was less prominent but phase-precession became stronger and spike theta-phases were more informative. This shift between rate and phase codes was regulated by a differential control of CA1 place cells by mainly CA3 or entorhinal inputs, gated by local interneurons. These findings suggest that environmental features influence the anatomical and computational substrate of hippocampal representation of space.

## RESULTS

### Superficial and deep CA1 place cells dominated in cue-poor and cue-rich environments, respectively

Dissecting the mechanisms governing place cells in freely moving animals across different environments is complicated due to the large number of uncontrolled variables such as head direction, physical interaction with objects, mixed contribution of distal and local cues (Knierim et al., 1998; Huxter et al., 2008; Ravassard et al., 2013; Jayakumar et al., 2019). To avoid such confounding factors, we tested head fixed mice (n = 6) running on a non-motorized treadmill to collect water rewards (Figure S1). The treadmill had a 230-cm belt with two zones, one without any local cue (‘cue-poor’ zone) and other enriched with several large tactile cues (‘cue-rich’ zone). Influence of distal visual cues was minimized by adding tall walls around the track. This task has the additional advantage that allows monitoring the same population of cells across both types of environments in each trial. We used high-density silicon probe electrodes to record the spiking of individual place cells and LFPs from the hippocampal CA1 region.

CA1 pyramidal neurons displayed place fields that tiled the whole treadmill belt and were stable throughout the recording session (Figure 1A and Figure S1). The incidence of place fields and their peak firing rates were similar in the cue-rich and cue-poor zones of the belt (Figure 1A-B; p > 0.05 Kruskal-Wallis test), although their width was smaller in the cue-rich zone (Figure 1C; p = 0.014). Mice ran with similar speeds in both zones (Figure S1, p > 0.05). We grouped place cells into cue-rich (n = 180) and cue-poor (n = 72) cells, according to the location of their peak firing. Next, we examined the anatomical distribution of the two groups of place cells. We defined the middle of the CA1 pyramidal layer as the recording site where sharp-wave ripples had the largest amplitude (Figure 1D; Mizuseki et al., 2011; Oliva et al., 2016b). A majority of the cells with place fields in the cue-rich zone of the belt were located above this point (deep sublayer), while those cells with fields in the cue-poor zone were preferentially located below the middle of the layer (superficial sublayer; Figure 1E-F; p = 0.003). Overall, 75.7% of all the place cells in the cue zone were deep pyramidal cells while 71.5% of place cells in the cue-poor zone were superficial pyramidal cells (Figure 1G). In addition, we found that cue-poor zone place cells had higher firing rates during sharp-wave ripples than cue-rich zone place cells (Figure S1, p = 0.021 Kruskal-Wallis test), in agreement with previous results that found higher within-ripple firing rates for superficial compared to deep CA1 pyramidal cells (Mizuseki et al., 2011; Valero et al., 2015; Oliva et al., 2016b). These results show that cue-rich and cue-poor environments are differentially represented in anatomically segregated populations of hippocampal place cells.

**Figure 1:**
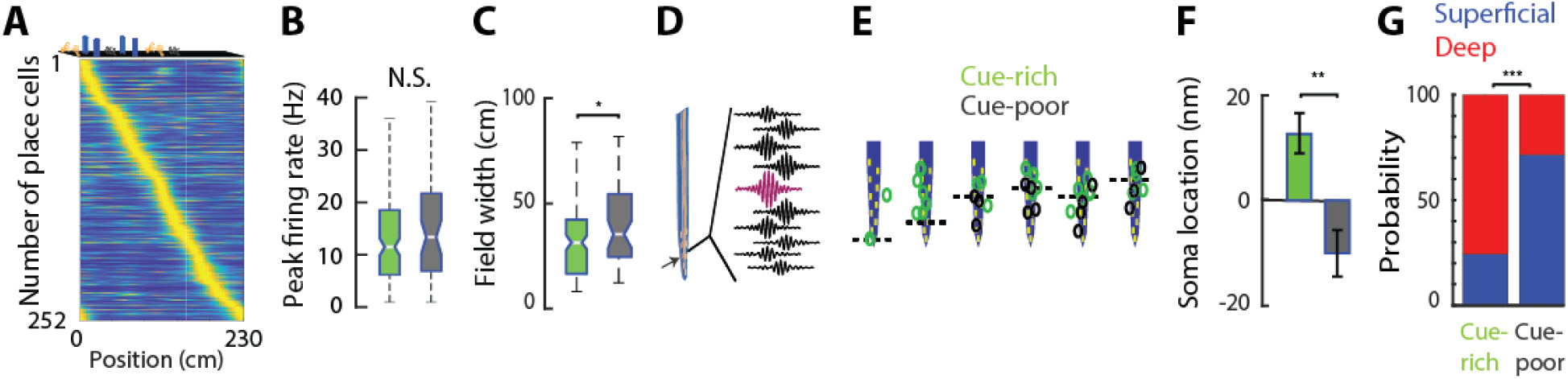
Segregated CA1 subpopulations for navigation in different environments. **A)** Plot showing firing maps of CA1 pyramidal cells on the belt (n = 252). Cells were sorted according to the location of their highest firing rate and rate maps normalized to this maximum value. Top: illustration of object distribution on the belt. **B**) Peak firing rate was not significantly different for cue-poor and cue-rich zone cells (p > 0.05, Kruskal-Wallis test). **C**) Place fields in the cue-poor zone were wider than those in the cue-rich zone (p = 0.014). **D**) Example filtered LFP traces (120-250 Hz) showing largest ripple amplitude in the middle of the CA1 pyramidal layer (magenta). **E**) Soma location of cue-poor (grey) and cue-rich (green) zone place cells relative to the middle of the pyramidal layer (dashed line) in an example animal. **F**) Mean ± SEM of soma location relative to middle of the pyramidal layer for all place cells (p = 0.003, one-way ANOVA). **G**) Fraction of place cells in the cue-rich and cue-poor zones in the deep or superficial CA1 sublayers, respectively (p = 9e-11, Fisher’s exact test).

### Different spatial codes in cue-poor and cue-rich environments

We asked whether hippocampal spatial representations of the two zones of the belt were supported by the same or different coding schemes. Firing rates of place cells in the cue-poor zone carried more spatial information than those in the cue-rich zone (Figure 2A,B; p = 0.014, Kruskal-Wallis test), were more spatially selective (Figure 2C; p = 0.017), less sparse (Figure S2; p = 0.038) and better spatially tuned (Figure S2; p = 0.027). Cue-rich zone place cells fired more spikes outside their place fields (Figure 2D; p = 0.019), which contributed to their poorer spatial rate code.

**Figure 2:**
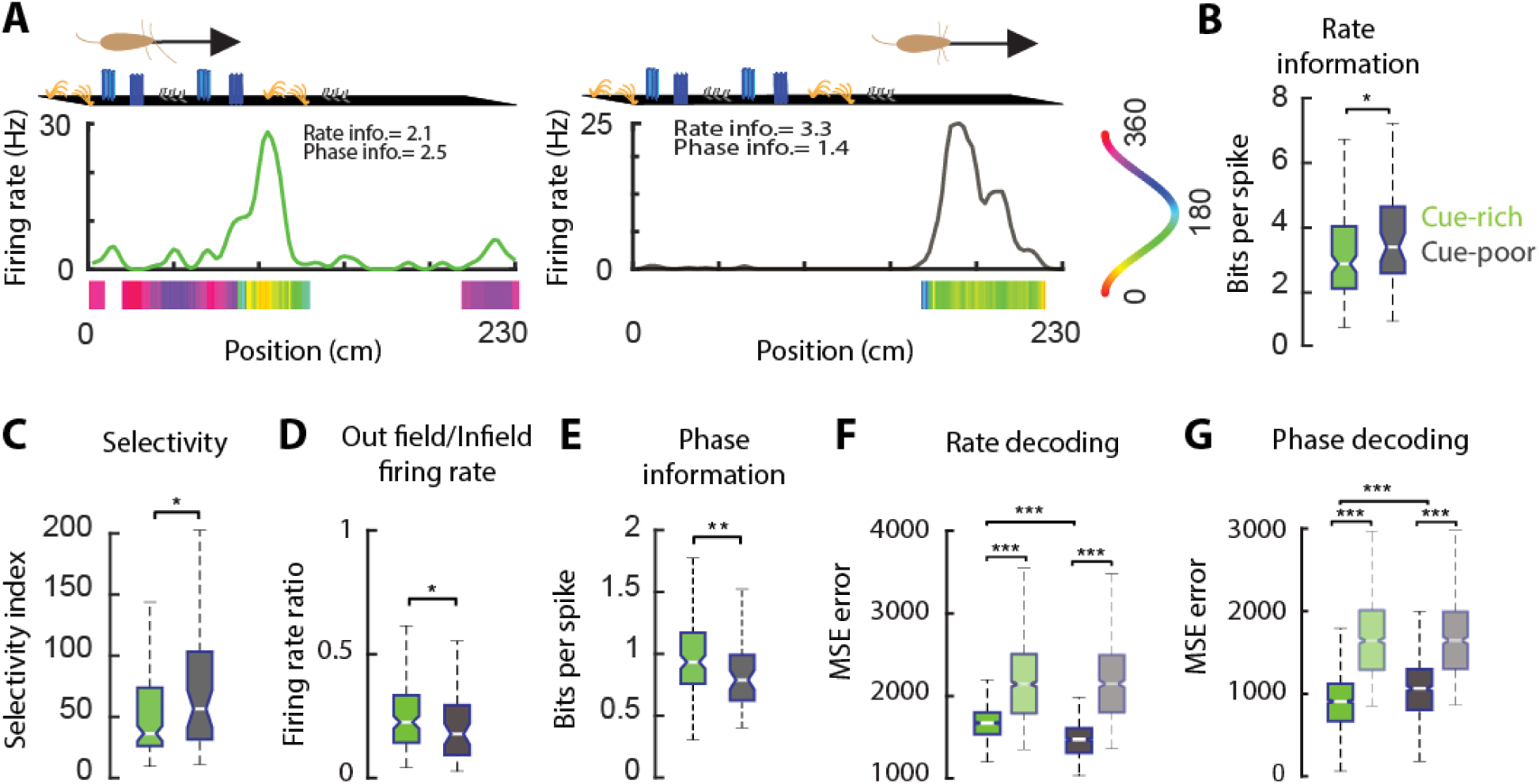
Spatial coding properties differ in the cue-rich and cue-poor zones of the belt. **A)** Example firing rate and spike-phase maps for one place cell in the cue-rich (left) and another in the cue-poo zone of the belt. Solid line is averaged firing rate per spatial bin across trials and color bar averaged spike theta-phase per bin. Arrow on the top indicate direction of running. **B**) Cue-poor zone place cells carried more spatial information in their firing rates and were more spatially selective (**C**) than cue-rich zone place cells. **D**) Cue-rich zone place cells fired more spikes outside their place fields than those in the cue-poor zone. **E**) Cue-rich zone place cells carried more spatial information in their spike theta-phases that cue-poor zone place cells. **F**) Spatial decoding accuracy with spike rates was higher in the cue-poor than in the cue-rich zone, while the opposite was true when using spike-phases (**G**). Lighter color boxplots are surrogate data distribution. */**/*** P < 0.05/ 0.01/ 0.001, Kruskal-Wallis test.

To assess the theta phase coding of spatial information, we calculated one-dimensional spike theta-phase maps the same way as rate maps but circularly averaging the instantaneous spike theta-phases instead of spikes rates. These maps revealed stereotyped phase patterns inside the place fields (Figure 2A and Figure S2). In contrast to the results obtained with firing rates, spike phases from cells in the cue-rich zone carried more spatial information than those in the cue-poor zone (Figure 2E; p = 0.0059, Kruskal-Wallis test).

To more directly compare how well the animal’s position was encoded by either the firing rates or spike phases of the same neuron, we employed a decoding approach. We trained a maximum correlation classifier to predict animal’s position using either the firing rate or spike-phases maps of individual neurons with 70% of the trials and compared the decoding error of predicting animal position in the remaining 30% of the trials, with the classifier trained with either firing rate or phase maps. We decoded position from both spike-rates and spike-phases with high accuracy (Figure 2F-G; p < 1e-67 for all cases when compared to randomly shuffled data; Kruskal-Wallis test). Mean square decoding errors using firing rates were significantly smaller for cue-poor zone place cells compared to cue-rich zone place cells (Figure 2F; p = 3.0e-88, Kruskal-Wallis test). In contrast, decoding errors with spike-phases were smaller for cue-rich zone place cells compared to those in the cue-poor zone (Figure 2G; p = 1.7e-15). These results suggest that the hippocampal rate code is more spatially informative in a cue impoverished environment but it degrades in a cue-rich environment, where the phase code becomes more spatially informative.

Place cells display a monotonic advance of their firing relative to the ongoing theta rhythm as the animal transverse the place field of the neuron, resulting in a systematic shift forward in successive theta cycles (‘phase-precession’; O’Keefe and Recce, 1993). We hypothesized that this phase-position correlation accounted for the strong predictive accuracy of animal position obtained with spike-phase maps. We thus examined phase-precession properties of place cells in the cue-rich and cue-poor zones of the belt (Figure 3 and Figure S2). We restricted this analysis to fields with significant phase-position correlation (n = 70 /45 cue-rich and cue-poor zone place fields; p < 0.05, linear-circular regression). Place cells in the cue-rich zone displayed steeper phase-precession slopes (Figure 3A-B; p = 0.0082) and spanned over a wider theta-phase range (Figure 3C; p = 0.0019) than those in the cue-poor zone. These differences can be explained by the fact that the phase of the spikes at the onset of the place field in cue-rich zone place cells were closer to the theta peak (360°) compared to those in the cue-poor zone (Figure 3A,D; p = 0.029). Place field width and peak firing rates were not significantly different between the subset of cue-rich and cue-poor zones place fields used for this analysis (p > 0.05 for both metrics), and, therefore, cannot account for the observed differences in phase-precession properties. The wider theta-phase range of the cue-rich zone place cells can also explain the higher spatial information and decoding accuracy obtained with spike-phases compared to those of cue-poor zone cells that only sampled a reduced phase space of the theta cycle.

**Figure 3:**
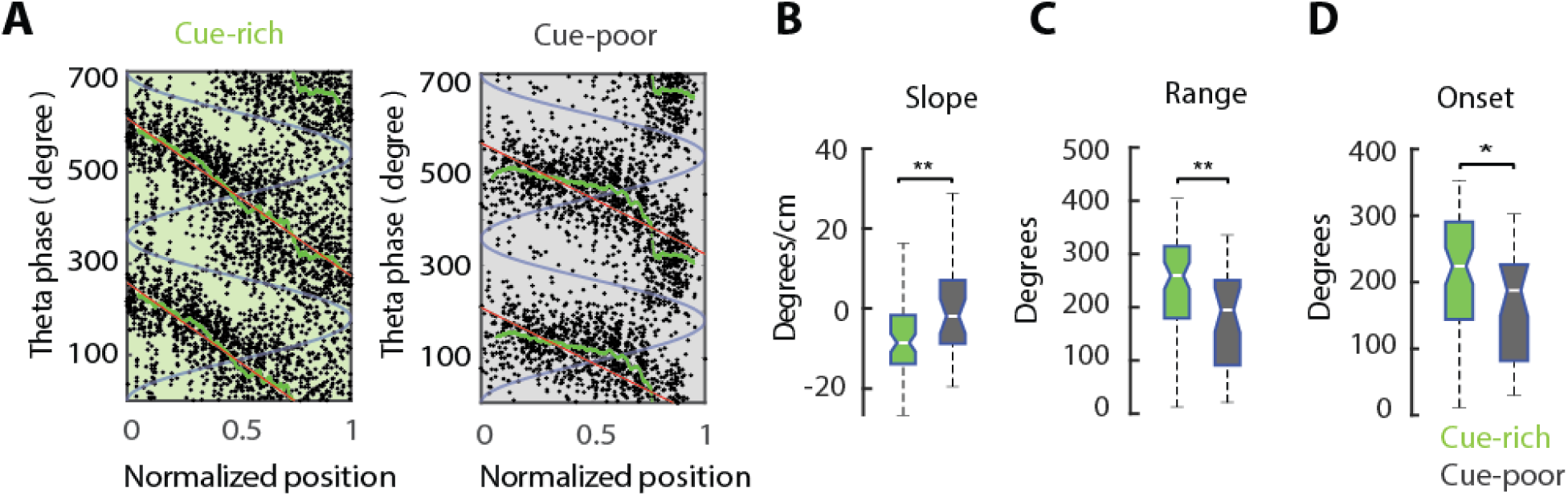
Wider phase precession in the cue-rich zone of the treadmill. **A)** Theta-phase versus normalized position inside the place field for two example place cells in the cue-rich (left) and cue-poor zones (right). Each black dot is one spike. Red line show phase-position correlation and green line mean theta phase. Two theta cycles (blue line) are shown for clarity. **B**) The slope of phase-precession was larger for place fields in the cue-rich zone than those in the cue-poor zone. **C**) Theta phase-range of spikes within the place field was larger in the cue-rich than in the cue-poor zone. **D**) Spike phases at the onset of the field were closer to the theta peak (360°) for place cells in the cue-rich zone. */** P < 0.05/ 0.01, Kruskal-Wallis test.

As a control, we changed the location of reward delivery and compared place field and spatial coding metrics between cue-rich and cue-poor zones. We found the same relationships (Figure S3), indicating that the observed differences in place cell properties were a consequence of the presence or absence of local cues on the belt rather than the location of the reward.

### Temporal compression of space by theta sequences reflects environment complexity

Our results from single cell spatial coding properties suggested that hippocampal representation of space has higher resolution on the cue-rich zone of the belt, compared to the cue-poor zone. We thus hypothesized that such differences in single neuron properties also alter representation of space by assemblies of hippocampal neurons.

As an animal runs across an environment, different place cell assemblies become successively active in a sequential manner along the travel trajectory. At any given location, the place fields that encode the current position overlap with the tails of adjacent place fields encoding previous and upcoming locations (Figure 4A). Place cells encoding past positions will fire earlier in the theta cycle, followed by those encoding current location firing at the theta through and those encoding future positions later in the cycle (Figure 4A). These “theta sequences” constitute temporally compressed ensemble representations of spatial trajectories (Skaggs et al., 1996; Dragoi and Buzsaki, 2006; Foster and Wilson, 2007). The degree of ‘temporal compression’ of space can be estimated as the ratio between the distance of two neighboring place fields peaks and the time lag between the spikes of the two corresponding place cells within one theta cycle (Figure 4B; Dragoi and Buzsaki, 2006).

**Figure 4:**
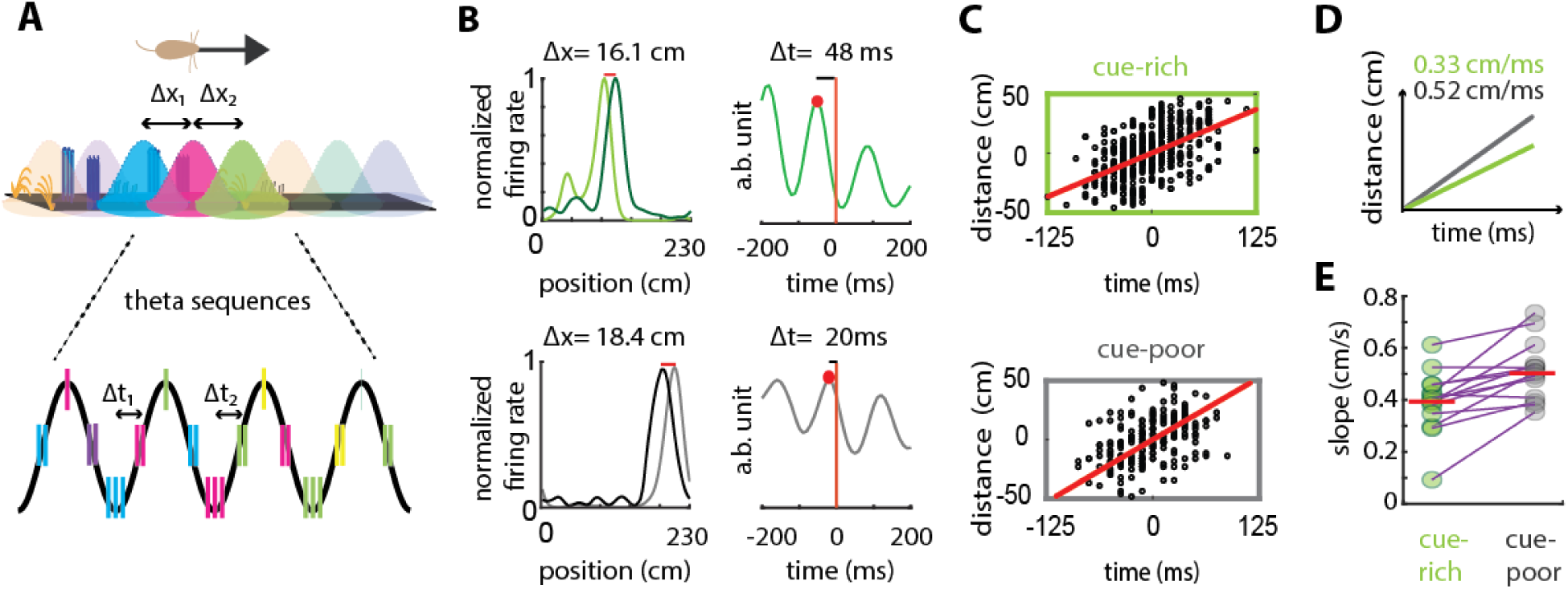
Environmental cues affect temporal compression of space by hippocampal theta sequences. **A)** Overlapping place fields along animal’s trajectory (top). Δx_1_/Δx_2_ = distance between the peaks of successive fields. Firing of example place cells (color of cell’s spikes match that of their place field) in successive theta cycles. Δt_1_/Δt_2_ = time lag between the firing of two place cells in the same theta cycle. **B**) Example of two overlapping place fields (left) and theta-scale cross-correlation of their spikes (right) for a pair of cue-rich (top) and cue-poor zone (bottom) place cells. Δx= distance between place field peaks; Δt= time lag between the two cell spikes in the first theta cycle (red dot). **C**) Correlation between place field peak distance and the theta time lag of their spikes for all overlapping CA1 place fields in the cue-rich (top; n = 611) and cue-poor (bottom; n = 216) zones. Red line indicates correlation slope (Δs/Δt). **D**) Compression slopes calculated in C) are displayed in the same axes for comparison purposes. **E**) Session by session comparison of compression slopes in the cue-rich versus the cue-poor zones (n = 13 sessions from 6 mice; p < 6.1e-4, t-test).

To examine whether the presence of local cues affected trajectory encoding by hippocampal theta sequences, we calculated the correlation between place field peaks and theta-scale time lags for all overlapping place field pairs in each zone of the belt (Figure 4C). As predicted from single neuron analysis, the slope of this correlation was significantly higher in the cue-poor zone than in the cue-rich zone (Figure 4D,E; 0.53 cm/ms vs 0.34 cm/ms; p < 6.1e-4, t-test), indicating a more compressed ensemble representation of space in the cue-poor zone. To verify that this difference in spatial compression was not due to changes in theta oscillation properties across zones, we found no differences of theta frequency, theta power or running speed between cue-rich and cue-poor zones (Figure S1, 4; p > 0.05, Kruskal-Wallis test). These results were maintained when reward location on the belt was changed (Figure S4)

### CA3 pyramidal cells have similar spatial coding properties to superficial CA1 cell

We found that superficial and deep CA1 cells expressed place fields preferentially in the cue-poor and cue-rich zones of the belt and that their spatial coding properties are manifested primarily as a rate or phase code, respectively. Because the CA3 input is the main driver of CA1 place cell firing (Csicvari et al., 2003; Dragoi et al., 2006; Nakashiba et al., 2008; Davoudi and Foster, 2019) and because CA3 input is more effective at driving superficial CA1 pyramidal cells (Valero et al., 2015; Masurkar et al., 2017), we nexy investigated spatial coding properties of CA3 neurons.

CA3 place fields were similarly expressed in both zones of the belt (Figure 5A and Figure S5; n = 149 cells in 6 mice). CA3 place cells in the cue-rich and cue-poor zones displayed similar differences in spatial coding properties as those of CA1; i.e., better rate coding in the cue-poor zone and better phase coding in the cue-rich (Figure 5B-H). Place fields in the cue-poor zone were wider than those in the cue-rich zone, also for CA3 (Figure S5; n = 40/109; p = 0.01, Kruskal-Wallis test). CA3 cells carried more spatial information in their firing rates compared to all CA1 neurons (Figure 5 B; p = 0.042/ 0.012 for cue-rich and cue-poor zones, Kruskal-Wallis test) and their fields were more spatially selective (Figure 5 C; p = 4.4e-5/ 0.024) and less sparse (Figure S5; p = 0.043/ 0.035). In contrast, CA3 spike-theta phases were less spatially informative than in CA1 (Figure 5D; p = 5.7e-31/ 0.023). We previously hypothesized that a lower spatial information of spike-theta phases could be the result of a reduced phase-precession range. In agreement with this hypothesis, the range and slope of CA3 phase precession were smaller than those of CA1 (Figure 5 E-F; p = 4.7e-6/0.023 for range and p = 0.018/0.028 for slope). Finally, we compared the accuracy of position decoding using CA3 firing rates or spikes phases. Decoding errors were smaller for CA3 than for CA1 cells when firing rates were used (Figure 5G; p = 1.8e-82/ 5.5e-4), but they were higher when spike-phases were used (Figure 5H; p = 0.023/ 1.7e-9). In summary, these results indicate that in CA3 rate coding is more reliable and phase coding is less reliable, compared to CA1. Consequently, CA3 spatial coding properties are more similar to those of CA1 superficial cells.

**Figure 5:**
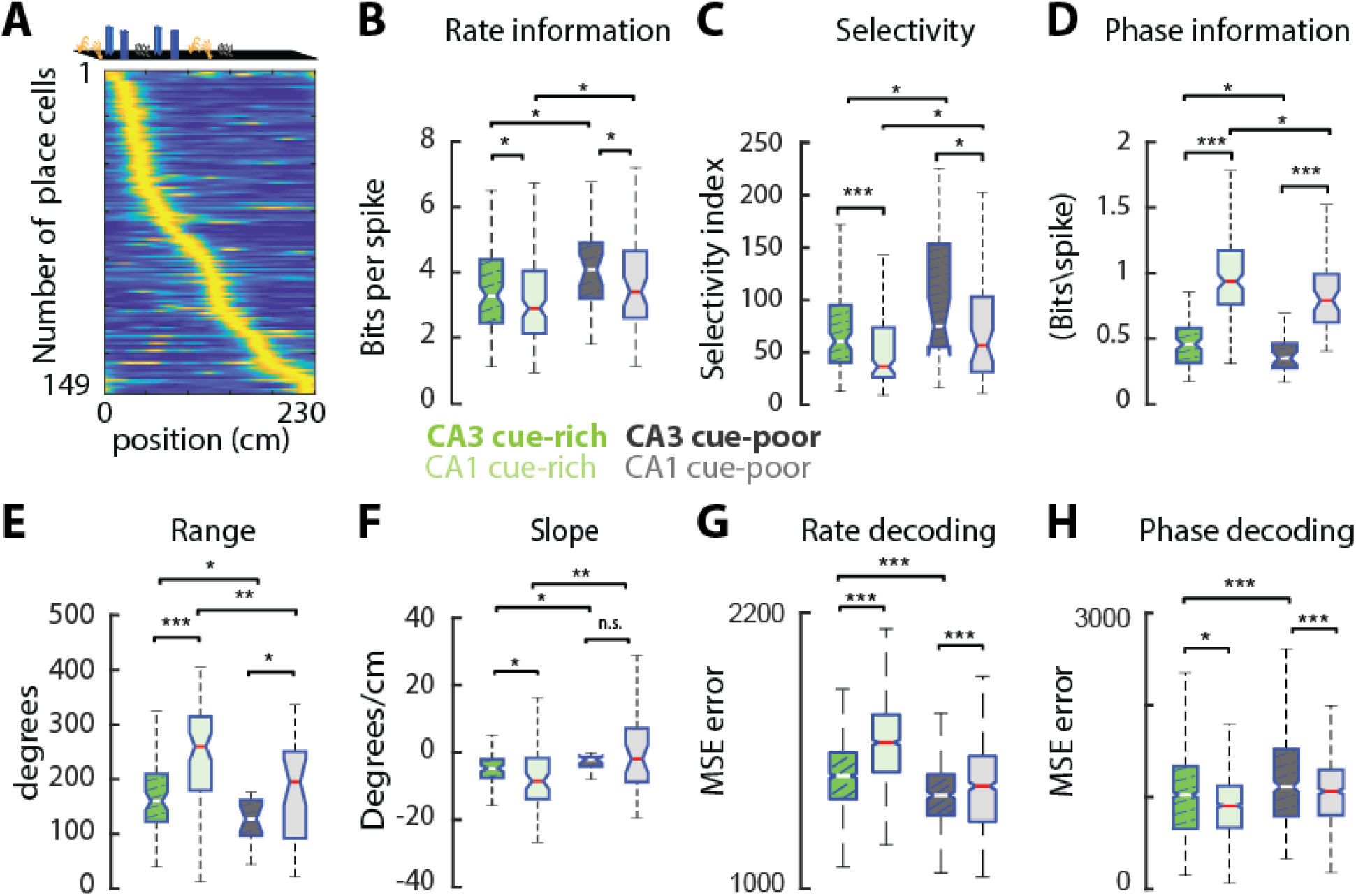
CA3 place cells have more precise spatial rate coding and poorer phase coding than CA1. **A)** Plot showing firing maps of CA3 pyramidal cells on the belt (n = 109 cue-rich and 40 cue-poor zone cells). Cells were sorted according to the location of their highest firing rate and rate maps normalized to this maximum value. **B**) CA3 place cells carried more spatial information in their firing rates than CA1 place cells. **C**) CA3 place fields were more spatially selective than those from CA1. **D**) CA3 place cells carried less spatial information in their spike-phases that CA1. **E**) Range and slope (**F**) of CA3 phase-precession were smaller than for CA1. **G**) Spatial decoding accuracy was higher in CA3 than CA1 using firing rates but lower using spike phases (**H**). Note that, for all metrics shown, trends between cue-poor and cue-rich zones were similar for CA3 than for CA1 place cells. */**/*** P < 0.05/ 0.01/ 0.001, Kruskal-Wallis test.

### CA3 and EC gamma inputs preferentially control place cells in cue-rich and cue-poor zones

Our previous results, together with existing anatomical evidence of stronger CA3 drive to superficial CA1 (Valero et al., 2015; Masurkar et al., 2017), suggests that stronger CA3 inputs in the cue-poor zone may contribute to the more efficient rate coding of CA1 in that zone. Yet, this mechanism alone cannot explain the wider phase-precession range and improved phase-coding in the cue-rich zone. Another key contributor can be the drive by layer 3 of entorhinal cortex (EC3), especially since the strength of EC3 input modulates the degree of phase-precession in CA1 (Schlesiger et a., 2015; Fernandez-Ruiz et al., 2017). Therefore, we compared the strength of CA3 and EC3 inputs to CA1 in the two zones of the belt.

We replicated previous results showing that these two inputs elicit gamma LFP oscillations of different frequency and theta-phase preference in CA1 (Figure 6A-B and Figure S6; Colgin et al., 2009; Schomburg et al., 2014; Lasztoczi and Klausberger, 2016; Fernandez-Ruiz et al., 2017; Lopes-dos Santos et al., 2018). We used the power of CA3 slow-gamma (30-50 Hz) and EC3 mid-gamma (60-100Hz) inputs to place cells in the cue-rich and cue-poor zones of the belt to estimate the strength of these respective inputs. Towards this goal, we detected mid- and slow-gamma frequency burst during running (Figure 6C). The power of CA1 mid-gamma events was larger in the cue-poor than in the cue-rich zone (Figure 6D; p < 2.4e-29, Kruskal-Wallis test). In contrast, the power of CA1 slow-gamma events (Figure 6E; p < 1.8e-100) and the ratio of mid-gamma to slow-gamma events (Figure 6 G, p = 1.4e-4, rank-sum test) were larger in the cue-rich zone. Slow-gamma power in the CA3 pyramidal layer was stronger in the cue-poor than in the cue-rich zone (Figure 6F; p = 2.1e-5, sign-rank test), mirroring CA1 slow-gamma.

**Figure 6:**
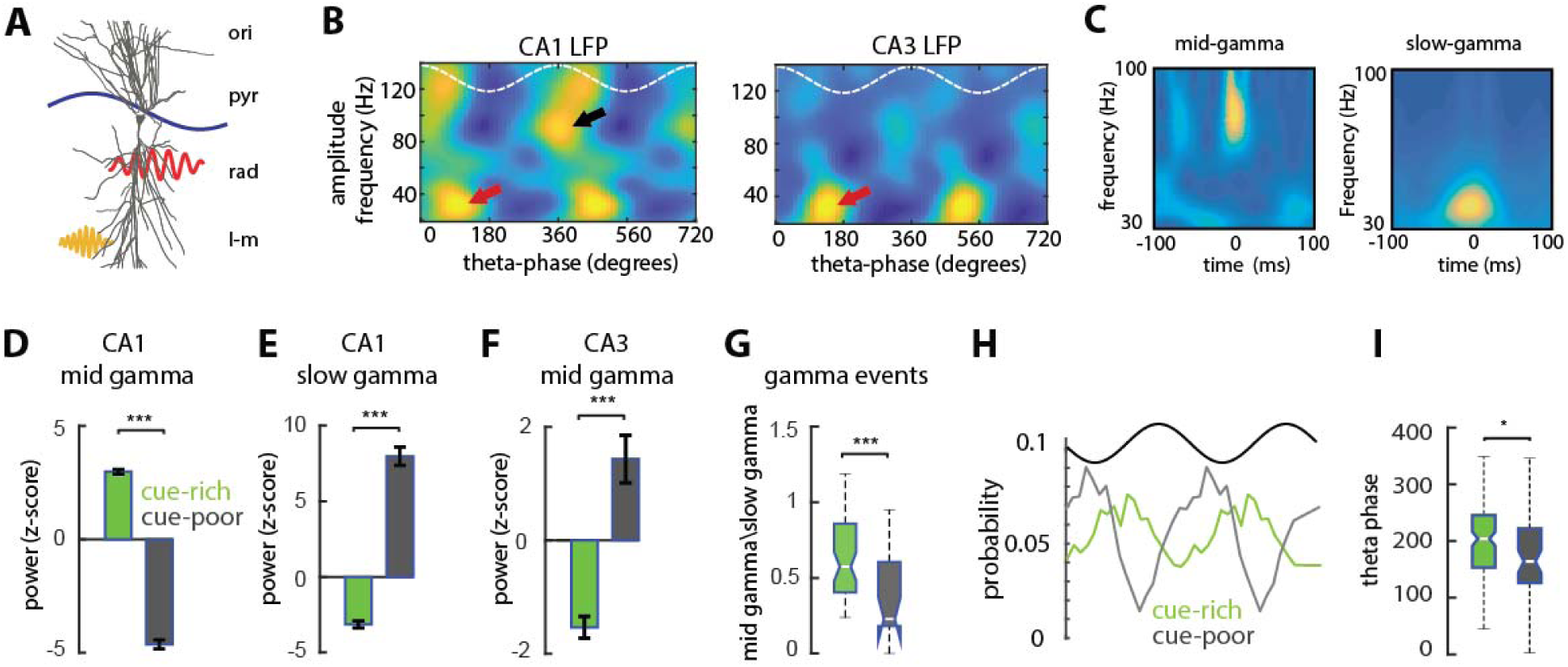
Low and mid gamma frequency inputs to CA1 place cells dominate in the cue-poor and cue-rich zones, respectively. **A**) Schema summarizing dendritic targets and temporal organization within the theta cycle (blue line) of the two main excitatory inputs to CA1 pyramidal cells: CA3 slow gamma (red) and entorhinal layer 3 (EC3) mid-gamma (yellow) inputs. ori= stratum oriens, pyr= pyramidal layer, rad= stratum radiatum, l-m= striatum lacunosum-moleculare. **B**) Average theta-phase gamma-amplitude comodulograms from CA1 (left) and CA3 (right) pyramidal layer LFPs (n = 6 mice). Two theta cycles (white dashed line) are show for clarity. Note the presence of two theta-modulated gamma components in CA1, slow (red arrow, 30-50Hz) and mid-frequency (black arrow, 60-100 Hz) with different theta phase preference and one in CA3 (red arrow, 30-40 Hz). **C**) Wavelet spectrograms of detected mid-gamma (left) and slow-gamma (right) bursts. **D**) Mean ± SEM CA1 mid-gamma and slow (**E**) power in the cue-rich and cue-poor zones displayed opposite trends (n = 7480 slow gamma and 4426 mid-gamma events in 6 mice). **F**) Ratio of occurrence of mid-gamma burst events compared to slow-gamma bursts in the cue-rich and cue-poor zones. **G**) CA3 slow gamma power was higher in the cue-poor zone, matching CA1 slow gamma distribution. **H)** Theta-phase distribution of firing probability for cue-poor and cue-rich CA1 cells. **I**) Preferred theta phase of firing for cue-poor and cue-rich cells.*/**/*** P < 0.05/ 0.01/ 0.001, Kruskal-Wallis test.

Since the relative strength of CA3 and EC3 inputs has been shown to modulate the theta-phase preference of CA1 spikes (Mizuseki et al., 2011; Fernandez-Ruiz et al., 2017; Oliva et al., 2016a; Navas-Olive et al., 2020), we hypothesized that, in the cue-rich zone, CA1 cells will fire closer to the theta peak, the preferred phase of EC3 input, while place cells in the cur-poor zone will fire closer to the theta trough, due to stronger CA3 drive. According to this expectation, cue-rich zone place cells fired closer to the theta peak than cue-poor zone place cells (Figure 6H-I; p = 0.012, Kruskal-Wallis test). These results suggest that CA3 input preferentially drive place cells in the cue-poor zone, while entorhinal inputs exert a stronger control over CA1 in the cue-rich zone. This interpretation is in agreement with a stronger entorhinal innervation of deep CA1 pyramidal cells (Masurkar et al., 2017).

### Recruitment of local interneurons is modulated by environmental features

Our findings so far indicate that the shifting strength of CA3 and EC3 inputs to CA1 place cells in the cue-poor and cue-rich zones of the belt drive the shift from a rate code-dominated to a phase code-dominated mode of spatial representation. Since the strength of interneuron recruitment by CA1 place cells is dynamically modulated by multiple factors such as learning, network synchrony or location within the place field (Dupret et al., 2013; Fernandez-Ruiz et al., 2017; Grienberger et al., 2017; English et al., 2017; McKenzie et al, 2019), we investigated whether short-term synaptic plasticity between CA1 pyramidal cells and local interneurons was also modulated as a function of environmental features and can thus contribute to the spatial coding differences we observed.

We identified putative monosynaptically connected pyramidal – interneuron cell pairs as determined by their cross-correlograms (English et al., 2017). The presence of a significant short-latency (1-3 ms) peak in the cross-correlogram denoted a functional monosynaptic cell pair (Figure 7A and Figure S7; n = 142 pairs in 6 mice). We compared spike-transmission probability for all cell pairs separating presynaptic spikes from the cue-rich and cue-poor zones and recalculating pyramidal cell – interneuron cross-correlograms in each zone (Figure 7A). Next, we compared the spike-transmission probability ratio as cue-rich spike transmission probability / cue-poor spike transmission probability. For individual cell pairs, spike-transmission probability ratio changed significantly between the cue-rich and cue-poor zones (Figure 7A,B; median change between zones = 0.74, p = 9.4e-6, sing-rank test). For those interneurons that had more than one presynaptic partner (n = 34), we found that the strength of spike-transmission probability across both zones followed the same pattern for all presynaptic partners in the same zone (Figure 7C-D and Figure S7; p = 8.2e-7, sing-rank test). In other words, if one of the pre-synaptic pyramidal cells had higher spike-transmission probability in the cue-poor than in the cue-rich zone, the other pre-synaptic partners were also more likely to drive more strongly the interneuron in the cue-poor zone (Figure 7C,D and Figure S7).

**Figure 7:**
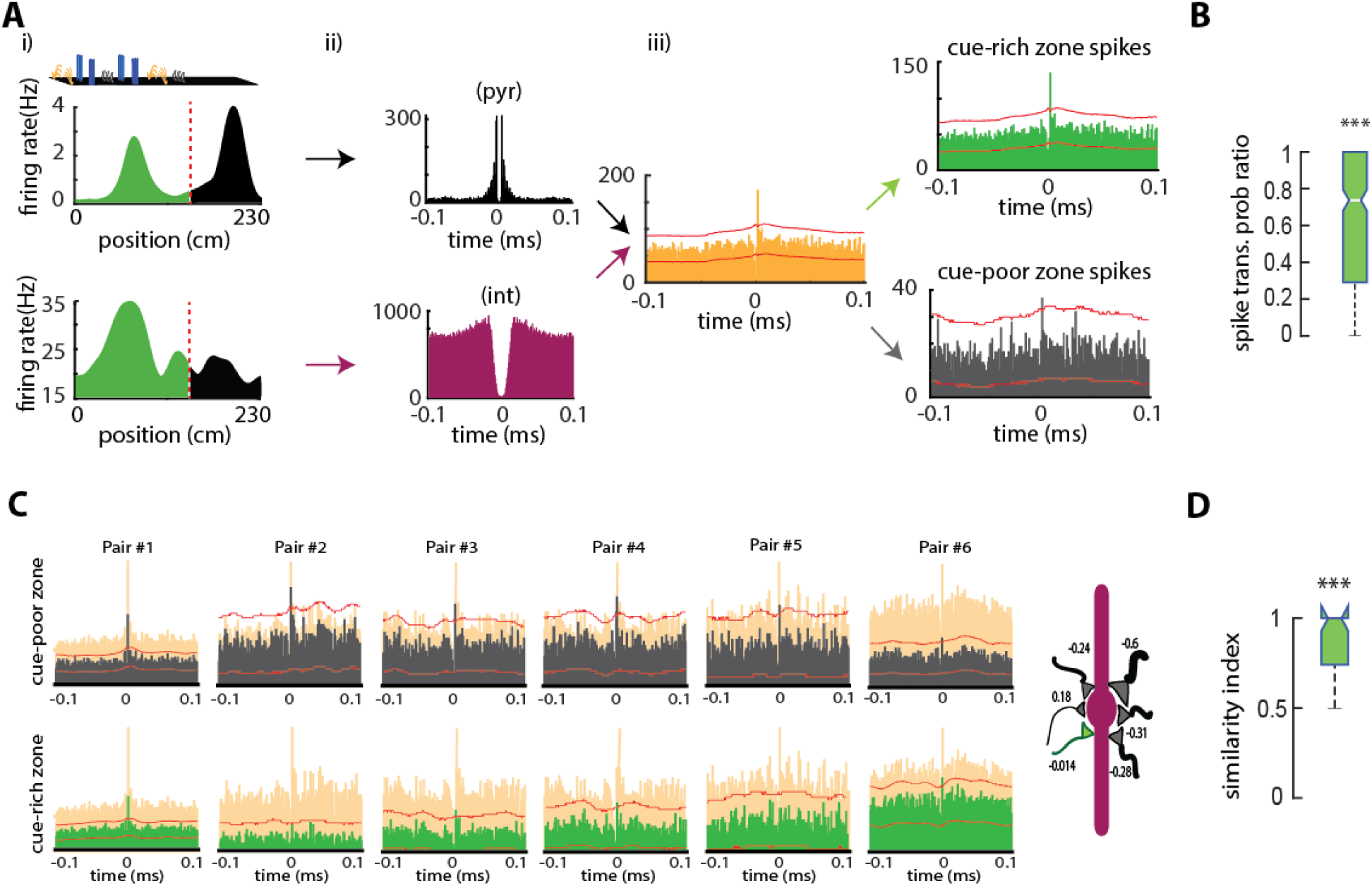
Variable recruitment of local inhibition in the cue-rich and cue-poor zones. **A)** Example monosynaptically connected CA1 pyramidal cell-interneuron pair. i) Firing maps for the pyramidal cell (top) and interneuron (bottom). The pyramidal cell has one place field in the cue-rich (green area) and another in the cue-poor zone (black). ii) On the left are auto-correlograms for the pyramidal cell (black) and interneuron (red). Spike trains’ cross-correlogram (in orange) displayed a sharp peak of firing probability for the interneuron 2 ms after the pyramidal cell fired. iii) Cross-correlograms constructed only from spikes in the cue-rich (green) or cue-poor zones (gray) indicate that the pyramidal cell only recruited the interneuron in the cue-rich zone. Red lines indicate significance levels (p < 0.001). **B**) Distribution of spike-transmission probability strength across the two zones of the belt (preferred zone - non-preferred zone/ (preferred zone + non-preferred zone)) for all significant pairs (n = 142) show a bias towards one of the two zones (p < 9.4e-6, sign-rank test). **C**) Example interneuron connected to six pre-synaptic pyramidal cells. Cross-correlograms for the cue-poor (black, top) and cue-rich (green, bottom) zone’s spikes are displayed relative to cross-correlograms with all spikes (orange). On the left, schema illustrating the relative strength (width) of each connection with the same interneuron. Note that, for 5 out of the 6 pairs, spike-transmission probability was stronger in the cue-poor zone (black synapses). **D**) Similarity index for all interneurons with more than one presynaptic pyramidal cell indicating a strong probability to share the same spatial preference for all the connections (n= 34; p < 8.2e-7, sign-rank test).

These results suggest that the spatial modulation of spike-transmission probability is modulated by environmental features. We also found that interneurons than were preferentially recruited in either the cue-poor or cue-rich zones differed in in their electrophysiological characteristics such as burstiness and recruitment during sharp-wave ripples (Figure S7), suggesting that different types of interneurons are recruited by active superficial and deep layer pyramidal cells.

### Place cell properties in 2D environments recapitulate the results during head-fixed running

Our controlled experimental preparation was effective to dissect the influence of local cues on hippocampal spatial coding. Yet, whether the results in head-fixed animals generalize to more natural behaviors and other species remained to be answer. To address this issue, we recorded rats (n = 5) while they were freely exploring either an open empty maze or the same maze with 4-6 large objects placed on it (‘objects maze’) to collect randomly scattered food rewards. We recorded place cells in both environments from CA1 and CA3 regions (Figure 8A-B; n = 249/185). In agreement with our observation in head-fixed mice, a larger proportion of deep CA1 place cells had place fields in the objects maze than in the open maze (n = 24 sessions; p = 0.02, rank-sum test), while the converse was true for superficial CA1 place cells (n = 24; p = 0.03).

**Figure 8:**
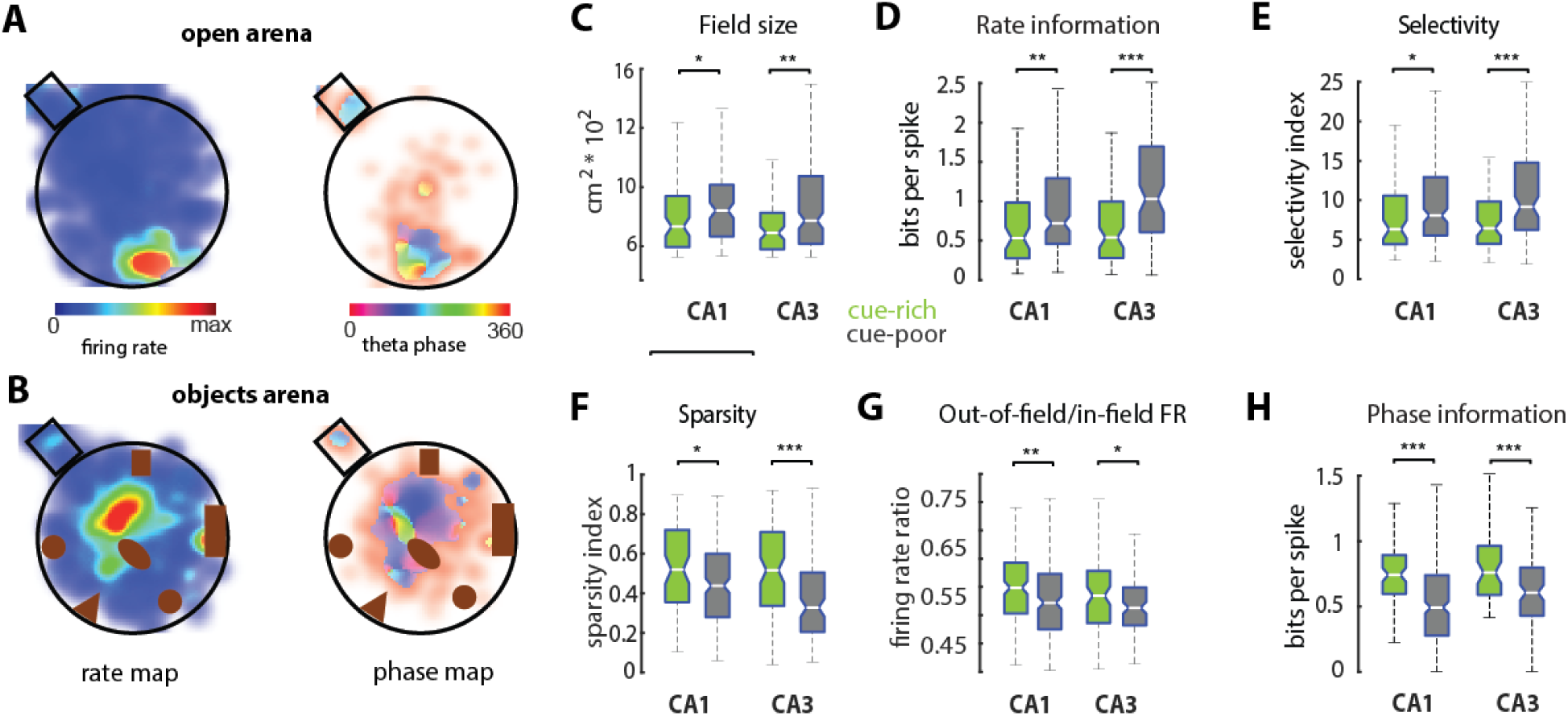
Rate and phase coding properties of place cells in 2D environments. **A**) Firing rate (left) and spike-phase (right) maps of representative CA1 place cells while a rat was exploring an open empty maze or the same maze with objects (**B**). **C**) Place fields were wider in the empty maze (n = 132/ 103 for CA1 and CA3) than in the objects maze (n = 117/ 82). **D**) Spatial information encoded in the firing rates of CA1 and CA3 place cells was higher in the open maze than in the objects maze. **E**) CA1 and CA3 place fields were more selective and less sparse (**F**) in the open maze than in the objects maze. **G**) Place cells fired more spikes outside their place fields in the objects maze than in the open maze. **H**) Spatial information encoded in the spike-phases of CA1 and CA3 place cells was higher in the objects maze than in the open maze. */**/*** P < 0.05/ 0.01/ 0.001, rank-sum test.

We calculated spike-rate and spike theta-phase maps in a similar way as before but in two-dimensions. Place fields in the objects maze were smaller than in the empty maze (Figure 8C; p = 0.03/0.003 for CA1 and CA3, rank-sum test) but their peak firing rates were not significantly different (Figure S8; p > 0.05). Firing rate of place cells in the open maze carried more spatial information than the firing of place cells in the objects maze (Figure 8D; p = 5.6e-3/ 4.7e-6). In addition, place cells in the open maze were more spatially selective (Figure 8E; p = 0.028 / 4.8e-4) and less sparse (Figure 8F; p = 0.016 /1.7e-5) than those in the objects maze. On the other hand, place cells in the objects maze fired more spikes outside their place fields (Figure 8G; p = 0.007/0.04). Spatial information content of 2-dimensional spike-phase maps in the open maze was lower than in the objects maze (Figure 8H; p = 5.8e-4/ 2.1e-3), in contrast to the result obtained with rate maps.

To verify that these differences in spatial coding properties were not due to non-specific changes in behavior or network activity, we compared animal velocity, spatial coverage, amount of exploration, theta power and frequency in both mazes but found no significant differences (Figure S8). In summary, we found consistent results in both head-fixed mice running on a treadmill and freely moving rats exploring a 2-dimensional maze. The spatial rate code in CA1 and CA3 place cells is less effective when the environment is rich in objects, whereas the timing of their spikes with respect to the theta rhythm is more spatially informative.

## Discussion

We addressed the question of how hippocampal circuits support navigation across environments differing in cue availability. We focused on two situations: feature-poor open environments in which a coarse low-resolution cognitive map is sufficient for navigation, and feature-rich environments where a finer-grain representation is needed due to the fragmented nature of the space. We observed differences in terms of cellular identity and computational mechanisms of the hippocampal spatial representation in these two types of environments. In cue-poor environments, a majority of active CA1 place cells were located in the superficial sublayer, while in cue-rich environments most of the recruited place cells resided in the deep sublayer of CA1 (Figure 1). These two subpopulations also favored different spatial coding mechanisms. Place cells in cue-poor environments had larger place fields with better spatial tuning than place cells in cue-rich environments. These differences resulted in a higher spatial information content and a more precise position decoding accuracy with firing rates of place cells in cue-poor environments (Figure 2). On the other hand, place cells in cue-rich environments had a wider phase precession, which resulted in an improved spatial encoding by their spike-theta phases compared to place cells in cue-poor environments (Figure 2 and 3). These differences were also reflected in the temporal encoding of trajectory representations by hippocampal assemblies, that were more compressed for cue-poor than cue-rich environments (Figure 4). The shift between a superficial CA1 rate coding-dominated mechanism to a deep CA1 phase code-dominated one was associated by a change in the excitatory and inhibitory inputs that control the firing of CA1 place cells (Figure 6 and 7). Finally, we demonstrated the universality of these results by replicating them in two different species (mice and rats) and in two different behavioral settings (head-fixed treadmill running and freely moving open maze exploration).

### Deep and superficial CA1 pyramidal cells utilize different spatial codes

Several recent studies have reported differences in place cell properties between deep and superficial CA1 pyramidal cells. Superficial place cells are more stable and discriminate better between contexts, while deep place cells are more modulated by reward and local landmarks (Mizuseki et al., 2011; Danielson et al., 2016; Geiller et al., 2017; Fattahi et al., 2018). Our present results extend these previous observations by showing that the hippocampal spatial map is preferentially encoded by superficial place cells in open environments, where a rate code dominates, and by deep place cells in those enriched with multiple local cues, where a phase code becomes more prominent. These findings suggest that deep and superficial cells express different codes to support efficient navigation across environments, providing the hippocampus with an enhanced computational flexibility. The coexistence of two specialized subpopulations with different spatial coding properties may allow the hippocampus to quickly adapt to changes in environmental features and to maintain the same time spatial representations at different resolution. Because deep and superficial pyramidal cells differ in their projection targets (Lee et al., 2014; Graves et al., 2016), their different spatial codes may allow different read-out mechanisms in downstream areas and serve to specialized functions.

The most striking differences in place coding properties have been described for CA1 pyramidal cells along the septo-temporal hippocampal axis. Ventral hippocampal place cells have larger fields and lower spatial selectivity than dorsal ones, suggesting that the ventral hippocampus encodes a lower resolution spatial map (Kjelstrup et al., 2008; Royer et al., 2010). These differences are reminiscent of our results for deep and superficial place cells. The fact that deep CA1 sublayer is largely expanded towards the ventral pole (Slomianka et al., 2011), possibly contributes to the observed differences in place coding. In addition, place cell properties also vary along the transverse (proximo-distal) hippocampal axis. Distal CA1 (closer to subiculum) place cells have multiple larger fields and are more strongly modulated by objects (Burke et al., 2011; Henrikssen et al., 2011; Oliva et al., 2016). Entorhinal cortex axons innervating distal CA1 originate on its lateral portion, where object coding cells are frequent, while those innervating proximal CA1 originate in the more medial part, where spatial coding is more prominent (Witter et al., 1989; Hargreaves et al., 2005). Taken together, these results suggest that spatial maps of different resolution are encoded simultaneously by segregated neuronal populations along the different anatomical axes. The fact that dorsal and ventral hippocampal cells strongly differ in their projection patters (Amaral and Witter, 1989; Witter et al., 1989) reinforces our interpretation that these spatial representations are conveyed to specialized postsynaptic “readers” and possibly serve to different behavioral functions.

### Excitatory and inhibitory inputs control the shift between rate and phase codes

Previous work suggested that the dynamic coordination between CA3 and entorhinal layer 3 gamma inputs control the spike timing of CA1 place cells (Bitter et al., 2015; Fernandez-Ruiz et al., 2017; Lasztoczi and Klausberger, 2016). In search for a mechanism that explains the shift between rate and phase codes across environments, we examined these inputs. We found that CA3 gamma input dominated in cue-poor mazes while entorhinal mid-gamma input was stronger in cue-rich environments (Figure 6). Anatomical and physiological data indicate a stronger entorhinal innervation of deep CA1 cells (Mizuseki et al., 2011; Masurkar et al., 2017), potentially explaining the increased proportion of active deep CA1 cells in cue-rich environments. It has been proposed that a stronger entorhinal input results in larger phase-precession range and slope (Schlesinger et al., 2016; Fernandez-Ruiz et al., 2017). Thus, a stronger entorhinal modulation of deep CA1 place cells can explain the wider phase-precession observed in cue-rich environments. At the population level, phase-precession is manifested as theta sequences, that encode spatial trajectories in a compressed manner (Skaggs et al., 1996; Dragoi et al., 2006; Foster and Wilson, 2007). This ensemble representation was more compressed in open environments than in cue-rich ones, reflecting a lower resolution spatial map (Figure 4). On the other hand, superficial CA1 pyramidal cells are more strongly driven by CA3 inputs (Valero et al., 2015; Masurkar et al., 2017). The present and previous data suggest that CA3 place cells are more selective and carry more spatial information in their firing rates than CA1 place cells (Leutgeb et al, 2004; Mizuseki et al., 2012; Oliva et al., 2016a). We hypothesize that rate-based spatial coding of superficial CA1 is inherited from the CA3 region and their reduced phase-precession is due to a weaker entorhinal drive compare to deep CA1 cells.

Local inhibition also plays an important role in modulating the firing of hippocampal place cells. Some models propose that place cells receive spatially uniform inhibition (Grienberger et al., 2017), while others suggest that different interneuron subtypes have specific influence on spatio-temporally tuned inhibition (Fernandez-Ruiz et al., 2017; McKenzie et al, 2019). Our finding that feedback inhibition onto CA1 place cells is strongly spatially modulated supports the later view (Figure 7). The finding that interneurons that were preferentially recruited in the cue-rich and cue-poor zones had different physiological properties suggest that different interneurons subtypes are specifically recruited by environmental features. Therefore, we hypothesize that the shift between rate and phase codes is gated by a change in the somato-dendritic distribution of inhibitory inputs that favor entorhinal distal dendritic depolarization versus CA3 proximal dendritic excitation.

Overall, our results suggest the co-operation of two subcircuits for spatial coding in the hippocampus. One involves mainly CA3-driven superficial CA1 neurons and favors rate coding. Another subcircuit is mainly controlled by entorhinal layer 3 targeting deep CA1 pyramidal neurons and favors phase coding. The shift between these two subcircuits may be supported by redistribution of perisomatic and distal dendritic inhibition.

### Behavioral relevance of the coexistence of multiple spatial coding mechanisms

What is the behavioral relevance of the coexistence of rate and phase codes for space in the hippocampus? In large open environments, an animal orients itself by distant cues and path integration. This can be achieved with a rate code, generated by the firing of individual place cells having an approximately gaussian distribution over a space of several tens of centimeters up to a few meters (Kjelstrup et al., 2008). At the population level, this rate code generates a coarse spatial representation that is sufficient to guide navigation between distant locations (O’Keefe and Nadel, 1978; Wilson and McNaughton, 1993; Leutgeb et al., 2004). Natural environments are often enriched of local landmarks and objects that offer additional reference points for navigation. In such situation, it is advantageous to have a high-resolution representation of the local environment. This may be better achieved with a phase code, since the phase of the spikes respect to the theta rhythm advances monotonically as the animal transverses the field of a place cell, conferring a spatial resolution of a few centimeters (Jensen and Lisman, 2000; Huxter et al., 2003; O’Keefe and Burgess, 2005; Tingley and Buzsaki, 2018). The ensemble of phase-precessing place cells generates a compressed representation of previous and upcoming positions in the form of theta-sequences (Skaggs et al., 1996; Dragoi and Buzsaki, 2006; Foster and Wilson, 2007). It has been proposed that theta-sequences are a mechanism that support prospective planning of behavioral trajectories (Johnson and Redish, 2007; Gupta et al., 2012). According to this hypothesis, a more compressed spatial representation (as we observed in the cue-poor zone of the belt) will allow for faster planning for travel in extended trajectories. On the other hand, a less compressed representation (as we found in the cue-rich zone) may only allow to plan shorter range trajectories but with higher spatial resolution. However, we need to emphasize that the dual-coding mechanism is complementary rather than exclusive. Contrary to common laboratory settings, in nature, environment complexity changes quickly, imposing different demands for animals to orientate and navigate. In order for the hippocampus to flexibly guide navigation, a mechanism to quickly switch between spatial maps of different scale is needed. We propose that the dynamic weighting of CA3 and entorhinal inputs to CA1 place cells drives the transition from a rate to a phase code-supporting circuit allowing the most advantageous spatial representation of changing environments.

## Author contributions

F.S., A.F-R. and S.R. designed the study and performed the experiments; F.S., A.F-R. and B.T. analyzed the data; A.F-R. and F.S. wrote the paper with input from the other authors.

## Acknowledgements

We thank Azahara Oliva, Kathryn Mcclain, Daniel Levenstein, Luke Sjulson, Ipshita Zutsi, Manuel Valero, Thomas Hainmueller and Peter Petersen for insightful comments on the manuscript and Alexander Lee for assistance with the experiments. This work was supported by a K99 grant (K99MH120343), NARSAD Young Investigator Grant and Sir Henry Wellcome Postdoctoral Fellowship (A. F-R); Korea Institute of Science and Technology Institutional Program (Project No. 2E30070; S.R.); R01 MH122391, U19NS104590, U19NS107616 (G.B.).

## STAR METHODS

### CONTACT FOR REAGENT AND RESOURCE SHARING

Further information and requests for reagents and resource may be directed to, and will be fulfilled by the Lead Contact, Dr. Antonio Fernandez-Ruiz.

### EXPERIMENTAL MODEL AND SUBJECT DETAILS

All experiments were conducted in accordance with institutional regulations (either Institutional Animal Care and Use Committee of the Korea Institute of Science and Technology or Institutional Animal Care and Use Committee at New York University Medical Center IACUC). 6 male C57BL/6 mice age between 6 and 7 weeks were used. The mice were housed by 2 to 3 per cage, in a vivarium with 12 hours light/dark cycles. Training and recording sessions described next occurred during the light cycles. Five male rats (Long-Evans, 3-5 months old) were used in this study. Rats were kept in the vivarium on a 12-hour light/ dark cycle and were housed 2-3 per cage before surgery and individually after it.

### METHOD DETAILS

#### Electrode Implantation and Surgery

Mice were put under isoflurane anesthesia (supplemented by subcutaneous injections of buprenorphine 0.1 mg/kg). Two small watch screws were driven into the bone above the cerebellum to serve as reference and ground electrodes. A plastic plate used for the head fixation on the treadmill was cemented to the skull with dental acrylic. After 3 weeks of training, the mice were put under isoflurane anesthesia and implanted with a 64-channel silicon probe (Neuronexus Buzsaki64) mounted on a micro-drive (Chung et al. 2017, Sariev et al. 2017). The silicon probe was slowly lowered into the hippocampus while monitoring electrophysiological activity and then retracted 200 µm above the pyramidal layer. The micro-drive was fixed on the skull and head-plate with dental cement. The craniotomy was covered with a bone wax and mineral oil mixture. A plastic cap was used to protect the micro-drive/silicon probe assembly. In order to let the electrode stabilize, the recordings were started at least three days after the implantation.

Rats were anesthetized with isoflurane anesthesia and craniotomies were performed under stereotaxic guidance. Animals were implanted with different types of silicon probes to record local field potential (LFP) and spikes across hippocampal regions and layers (Fernandez-Ruiz et al., 2017; Fernandez-Ruiz et al., 2019). Probes were mounted on custom-made micro-drives to allow their precise vertical movement after implantation. Probes were inserted above the target region and the micro-drives were attached to the skull with dental cement. Craniotomies were sealed with sterile wax. Two stainless steel screws were driven bilaterally over the cerebellum to serve as ground and reference for the recordings. Several additional screws were driven into the skull and covered with dental cement to strengthen the implant. Finally, a copper mesh was attached to the skull with dental cement and connected to the ground screw to act as a Faraday cage, attenuating the contamination of the recordings by environmental electric noise. After post-surgery recovery, probes were moved gradually in 50 to 150 μm steps per day until the desired position was reached. The pyramidal layer of the CA1 region was identified physiologically by increased unit activity and characteristic LFP patterns (Ylinen et al., 1995; Oliva et al., 2016). The identification of dendritic sublayers was achieved by the application of CSD and ICA analysis to the LFPs (Fernandez-Ruiz et al., 2012; Fernandez-Ruiz et al., 2013; Schomburg et al., 2014).

#### Head-fixed treadmill experiment

The treadmill was not motorized and consisted of a long velvet belt laying on 2 plastic 3D-printed wheels. Two pairs of LED and photo sensors read a disk pattern and measured forward and backward movements while another LED/photo sensor pair detected a hole on the treadmill belt and implemented the zero position. An Arduino board (Arduino Uno, arduino.cc) received these signals, computed the mice position in real time and controlled the reward delivery valves. Mice position, time and reward, was transmitted from the Arduino board to a computer via USB serial communication.

A 250-channels recording system (Intan Technologies, RHD2132 amplifier board with RHD2000 USB Interface Board and custom-made LabView interface) was used for acquiring continuously both the wideband neurophysiological signals and the treadmill signals at 30000Hz.

Mice (n = 6) were put under a water restriction scheme (1 ml per day) and trained to run for water reward on the treadmill. Specifically, the mice ran to receive water rewards that were delivered via a lick port on each trial at the same location of the belt. The water was sucked out of the lick port 15 cm after the reward delivery such that mice had to stop in that 15-cm section of the belt to consume the reward. The trials (complete rotation of the belt) started and ended when the mouse crossed the reward delivery point. Mice typically completed >18 trials per session. We recorded CA1 during 4 days. Then we lowered the shanks in CA3 and recorded CA3 for 5 days; overall, recordings were performed over 9 consecutive days.

Each belt (Figure S1) was 230-cm long and 5-cm wide and divided into two sections, one enriched with cues and the other without cues. The cues consisted of 3 pairs of identical erected objects (overall 6 objects) made of fuzzy rubbery wires (2-cm-high and 5-cm-long), horizontally cut shrink tubes (2-cm-high and 5-cm-long) and plastic dish pieces (1-cm-high and 10-cm-long), and were fixed on both sides of the belt to provide visual and tactile stimulation without impeding the mouse movement. The cue-rich zone spanned from the beginning of the first object to the end of the final object (0 to 150 cm) while the rest of the belt (150 cm to 230 cm) was defined as the cue-poor zone.

#### Freely moving open arena exploration

After surgery, animals where handled daily and accommodated to the experimenter, recording room and cables for 1 week before the start of the experiments. Prior to the start of the experiment animals were food restricted. One 30 to 60-minute-long behavior session was conducted daily, preceded and followed by 1 to 3 hour-long sleep sessions. The task consisted on free exploration on an elevated open circular arena without walls (150 cm diameter). To encourage animal exploration, small cereal crumbles were randomly scattered by the experimenter. The session was ended when the animal stop exploring for more than two minutes.

#### Histology

On the last day of recording, the animals were deeply anesthetized and transcardially perfused with 4% paraformaldehyde in phosphate buffer. The brain was removed and kept overnight in 4% paraformaldehyde solution. Coronal sections (100 µm thick) were obtained using a vibratome and mounted on slides using mounting medium with DAPI (Vector Laboratories, Inc). Images of DAPI and DiI fluorescence were acquired separately with a Nikon FN1 microscope equipped for fluorescence imaging.

#### Spike Sorting and Unit Classification

The wideband signals were digitally high-pass filtered (0.8-5 kHz) offline for spike detection or low-pass filtered (0-500 Hz) and down sampled to 1000 Hz for local field potentials. Spike sorting was performed semi-automatically, using KlustaKwik (klustakwik.sourceforge.net; Kadir, et al., 2014), followed by manual adjustment of the clusters with Klusters (Hazan et al., 2006) We recorded a total of 2084 neurons (1450 from CA1; 636 from CA3) during 9 recording sessions (CA1, 4; CA3, 5) following standard criteria for unit detection and clustering (Schmitzer-Torbert, et al. 2005, Hazan, et al. 2006, Kadir, et al. 2014).

Place cells were classify by zone of the belt encoded: if the peak firing rate of a place cell occurred in the cue-rich area, the cell was assigned to the cue-rich zone cell group, and if the peak firing rate occurred in the cue-poor zone area, the place cell was assigned to the cue-poor zone cell group.

To determine the cells’ position relative to deep and superficial CA1, we aligned and averaged all the ripples detected by a shank during the whole recording session and used the channel with the largest signal as a reference for the position of each cell (Figure 1 D-G). Spikes of all ripple episodes were combined to estimate the cells’ firing rate during the ripples (Figure S1 D).

#### Place field identification and metrics

To calculate firing rate maps, we divided the length of the belt into 100 pixels, divided the number of spikes in each pixel by the time the animal spent in each pixel (Figure S 1B) and then averaged across all trials (Figure 1 A). Only periods in which the animals run with a velocity higher than 5 cm/s were included.

The width of a place field was defined as the number of consecutive bins for which the mean firing rate was greater or equal to 10% of the peak firing rate (Figure 1 C).

The instant speed of the animal was calculated by dividing the time spent in each pixel by the size of the pixel (2.3 cm) (Figure S1 C).

Infield and out-field firing rates were calculated by averaging the pixels inside and outside the edges of the fields, respectively (with the edges defined as the positions where the mean firing rate reaches 10% of the peak firing rate).

The place fields tunity (Figure S2 B) was defined as the mean vector length of spikes position, after converting the belt position into circular coordinates. The analysis included cells that passed the Rayleigh test (p< 0.01).

Spatial information content (Skaggs et al., 1993) was calculated in bit per spike as following:

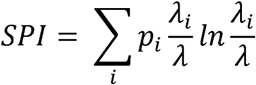

where *λ*_*i*_ is the mean firing rate of a unit in the i-th bin, *λ*_*i*_ is overall mean firing rate and *p*_*i*_ is the probability that animal being in the i-th bin (occupancy in the i-th bin / total recording time).

Place fields in 2D and corresponding metrics were calculated in an analogous manner.

#### Phase analysis

To analyze phase precession, LFP from the CA1 pyramidal layer was bandpass filtered in range of [6 12] hertz and the instantaneous theta phases of spikes was computed. We used (Kempter et al, 2012) method to quantify circular-linear association parameters such as onset, range, and slope of the phase precession. Spike phases in the range of [0 720] degree and their instantaneous position was feed to the algorithm. In addition, range of the slopes are restricted to a useful range by using optimization algorithm such as genetic algorithm the best slope will be selected among them. At the end, only those cells that pass criteria of having circular-linear phase precession (p-value of the circular-linear correlation coefficient <0.05) were selected for the analysis.

Phase maps of place cells were calculated the same manner as rate maps but using spike mean phase instead of spike count in each pixel.

To quantify the degree of temporal compression, we first detected overlapping place fields by computing for all cell pairs the cross-correlograms of spike times. Two neurons’ place fields overlapped if their cross-correlogram showed at least one local maxima matching a theta cycle (125 ms) and if the amplitude of the local maxima exceed 5% of the cross-correlogram peak amplitude. The theta time lag between the two cells was the time lag of the first local maxima (starting from 0). Next, we divided the distance of the place fields peaks by the theta time lag to estimate the spatial compression index. (Figure 4B; Dragoi and Buzsaki, 2006). To calculate the slope of the fitted line to the spatial compression indices, we used linear regression method from the supervised learning of Matlab fminunc function. (Figure 4 C-D)

#### Decoding analysis

Rate and phase maps were generated as described above and feed to the maximum correlation coefficient classifier (Meyers, 2013). The trained classifier generates a template for each class by averaging all training data point. After that, Pearson’s correlation coefficient between the points of each test class and template is calculated and maximum correlation value will be chosen as the predicted label. For each neuron, we used 70% of trials of rate or phase map to train the classifier and 30% of the trials were used for testing and predicting animal position. Performance of the model was quantified by calculating the mean squared error between the label and predicted position. To get more consistent predictive error we did at least 10-fold cross validation with different randomly selected 70 and 30 percent training and test trials. To confirm that model performance is not achieved by chance we trained the classifier by shuffled data (randomly permuting bins) and generated chance MSE values. Then we compare the real with the chances MSE values.

#### Gamma LFP analysis

Periods of slow and fast gamma activity in the LFP were calculated as described before (Colgin et al., 2009, Bieri et al., 2014). For both slow and fast gamma frequency bands, time points where the power was 2 standard deviation above the mean power were collected and used as reference for 200-ms time windows (Figure 6 C). Repeated time points or overlapping time windows (windows closer than 100 ms) across the frequency bands were discarded.

For each channel where cue-poor or cue-rich zone cells were recorded, slow and mid gamma episodes were detected during running states as described above. Individual power spectra were computed for each episode and normalized by the sum of their pixels. Then power spectra averages (across episodes) were computed for slow and mid gamma episodes.

#### Analysis of monosynaptic cell pairs

Cross-correlation (CCG) analysis has been applied to detect putative monosynaptic connections (Bartho et al., 2004; English et al., 2017). CCG was calculated as the time resolved distribution of spike transmission probability between a reference spike train and a temporally shifting target spike train. A window interval of [-5, +5] ms with a 1-ms bin size was used for detecting sharp peaks or troughs, as identifiers of putative monosynaptic connections. Significantly correlated cell pairs were identified using a previously ground-truth validated convolution method (English et al., 2017). The reference cell of a pair was considered to have an excitatory monosynaptic connection with the refereed neuron, if any of its CCG bins within a window of 0.5-3 ms reached above confidence intervals. To calculate similarity index, first we classified interneurons that had more than two presynaptic partner (n = 34), into two subgroups: group one were interneurons which receive stronger pre-presynaptic input from cue-rich zone spikes and group two were interneurons which receive stronger pre-synaptic input from cue-poor zone spikes. For example, in (Figure 7C) we showed an interneuron that was receiving 6 pre-synaptic input. For each pair, transparent orange cross-correlogram in the background shows transmission probability from all spikes and green or gray cross-correlograms show transmission probability from cue-rich or cue-poor spikes, receptively. To assign each pair as a cue-rich or cue-poor zone pair, we calculated the pair index as:

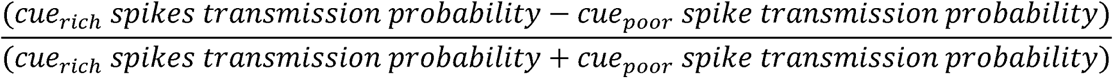

thus, the interneuron depicted in (Figure 7C) was receiving majority of its pre-synaptic input (5 out of 6) from cue-poor zone spikes (schema depicted in bottom right of Figure 7C). After calculating all the pair indices, we are able to check if the excited interneuron was preferentially recruited by cue-poor or cue-rich zones spikes via defining a similarity index as:

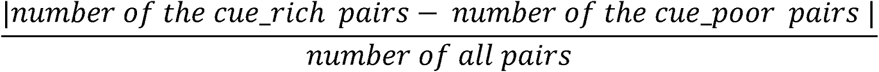

for example, the similarity index for interneuron in figure 7C was 0.83.

### QUANTIFICATIONS AND STATISTICAL ANALYSIS

All statistical analyses were performed in Matlab (MathWorks).

Kolmogorov-Smirnov test was used to determine if sample distribution was standard normal distribution. If normality was uncertain, nonparametric tests were used as described in following.

One-way ANOVA was used to determine whether different groups of an independent variable have common mean. To check whether data in each group has the same distribution Kruskal-Wallis one-way analysis of variance were used. Box-plots represent median and 25th 75th percentiles and their whiskers the data range. In some of the plots outlier values were not represented but they were always included in the statistical analysis.

Circular Statistics Toolbox for Matlab (Berens 2009) was used for comparison in polar coordinates. Nonparametric multi-sample test for equal medians similar to a Kruskal-Wallis test was used for linear data to determine whether population have common distribution. To check whether population is uniformly distributed around the circle Rayleigh test was applied. V test was used to determine whether there is a specific mean direction in each data set.

### DATA AND SOFTWARE AVAILABILITY

Part of the dataset included in this study is already available in the CRCNS.org and in the buzsakilab.com databases. The rest is currently under preparation to be deposited in the same databases but will be immediately available upon reasonable request.

Custom Matlab scripts can be downloaded from https://github.com/buzskilab/buzcode.

## Supplemental Figures and legends

**Figure S1.**
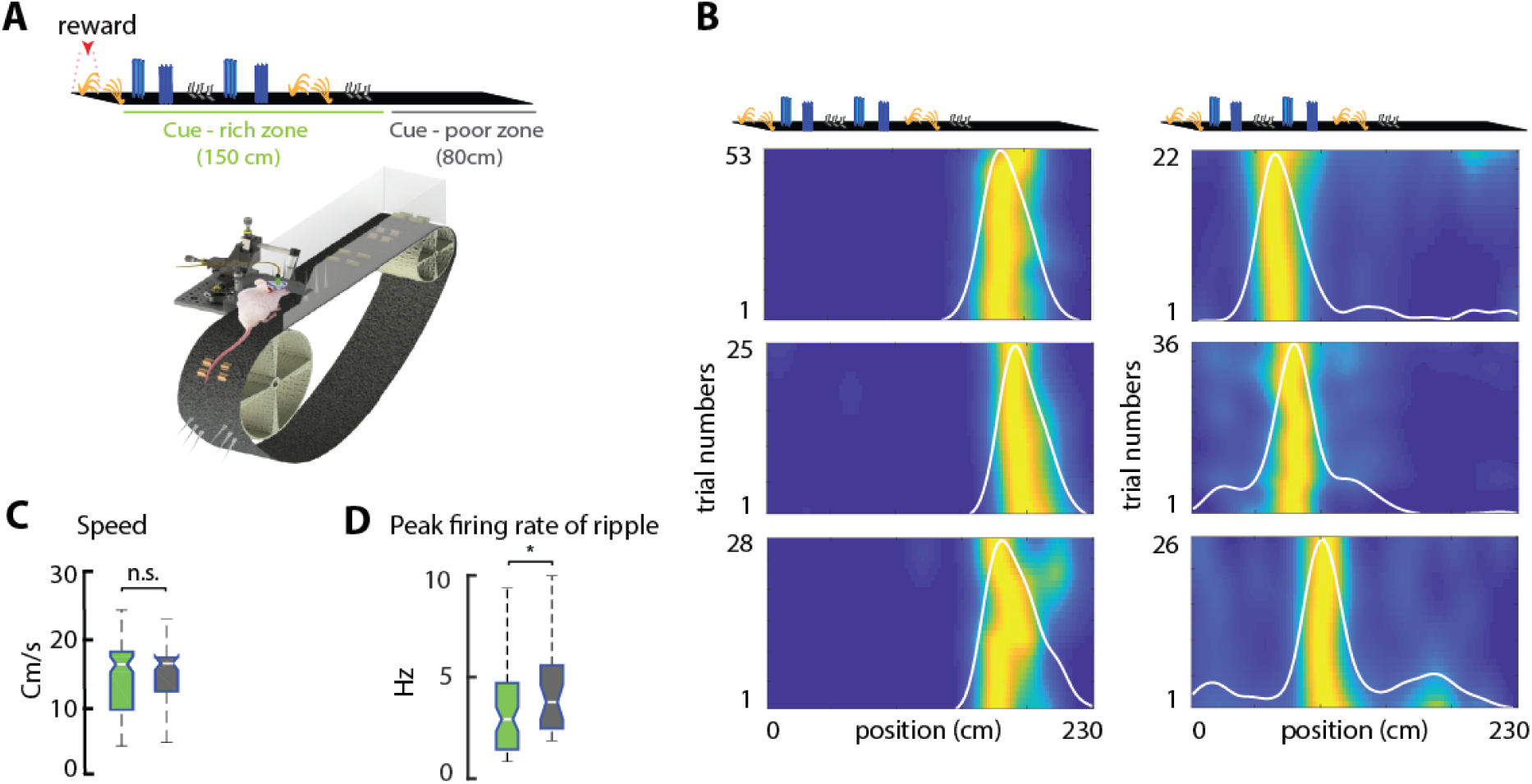
Place cell recordings in the treadmill. Related to Figure 1. **A)** Schematic of the treadmill setup for head-fixed mice. The treadmill was equipped with a 230 cm long belt displaying a zone enriched with multiple visual-tactile cues (‘cue-rich zone’) and other without any cues (‘cue-poor zone’). Walls around the track prevent the mouse to see distal visual landmarks. **B)** Examples of CA1 place cells recorded in the treadmill showing their stability across trials. White line is averaged firing rate (normalized) for the session. Color-coded map, smoothed trial-by-trial firing rate. **C)** Average running speed in the cue-rich and cue-poor zones across all sessions showed no significant differences (p > 0.05, Kruskal-Wallis test). **D)** Firing rate during sharp-wave ripples was higher for cue-poor zone than for cue-rich zone place cells (p = 0.021, Kruskal-Wallis test).

**Figure S2.**
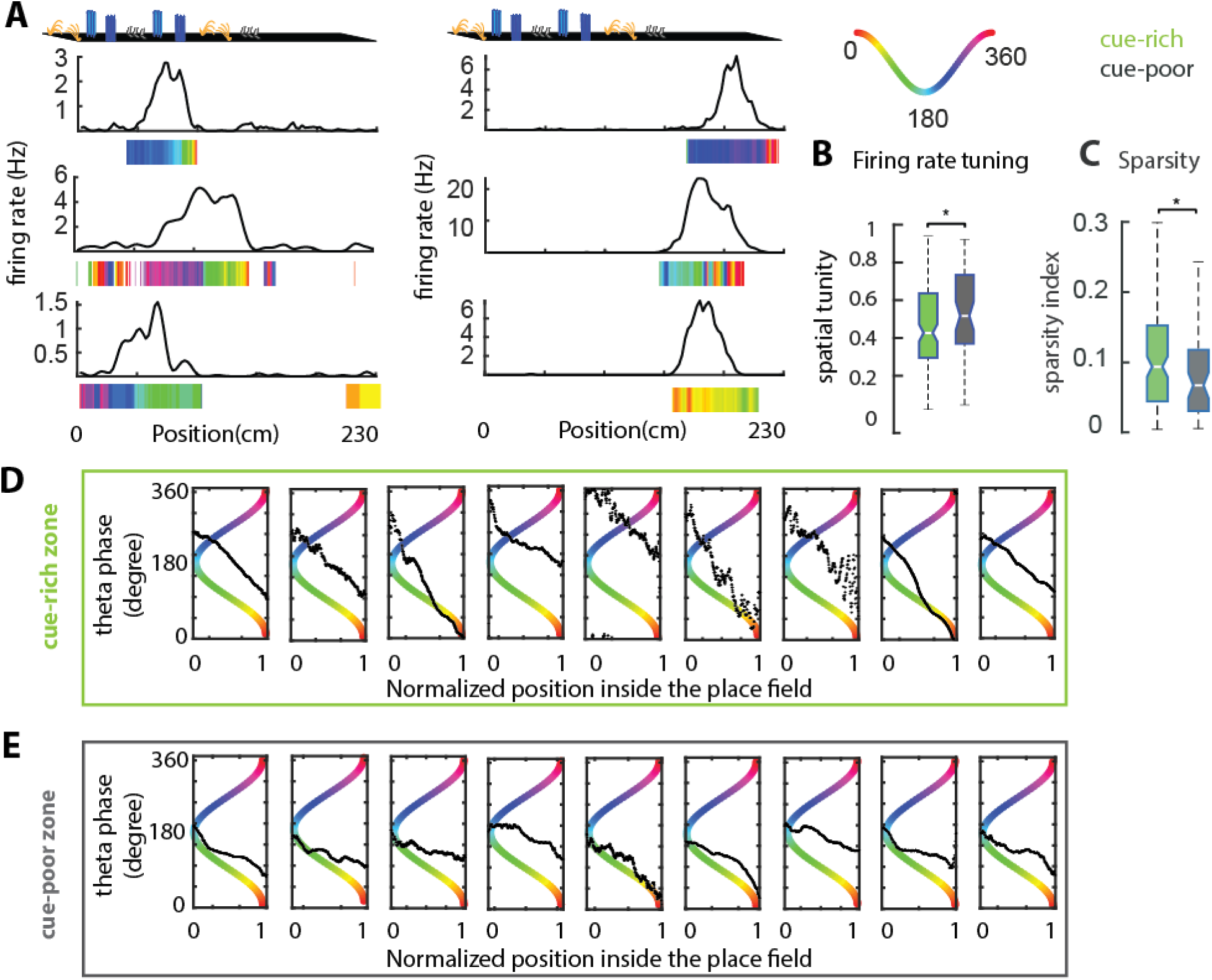
Firing rate and phase maps of place cells in the treadmill. Related to Figure 2 and 3. **A)** Additional examples of firing rate curves (black lines) and phase maps (color bars) for cue-rich zone place cells (left) and cue-poor zone place cells (right). **B**) Cue-poor zone place fields had higher firing rate spatial tuning than cue-rich zone place fields (p = 0.023, Kruskal-Wallis test). **C**) Cue-rich zone place cells had sparser firing maps compared to cue-poor zone place cells (p = 0.038). **D)** Examples of phase precession chosen from different sessions and animals for cue-rich zone place cells (first row) and (**E**) cue-poor zone place cells, (second row). Place fields were normalized from 0 to 1 and, in all cases, mice were running from left to right. Black lines are average theta phase of place cell spikes inside the field. Note the wider range of phase precession in the cue-rich zone and the spike-phase closer to the theta peak (360°) at the onset of the field, resulting in a steeper slope

**Figure S3.**
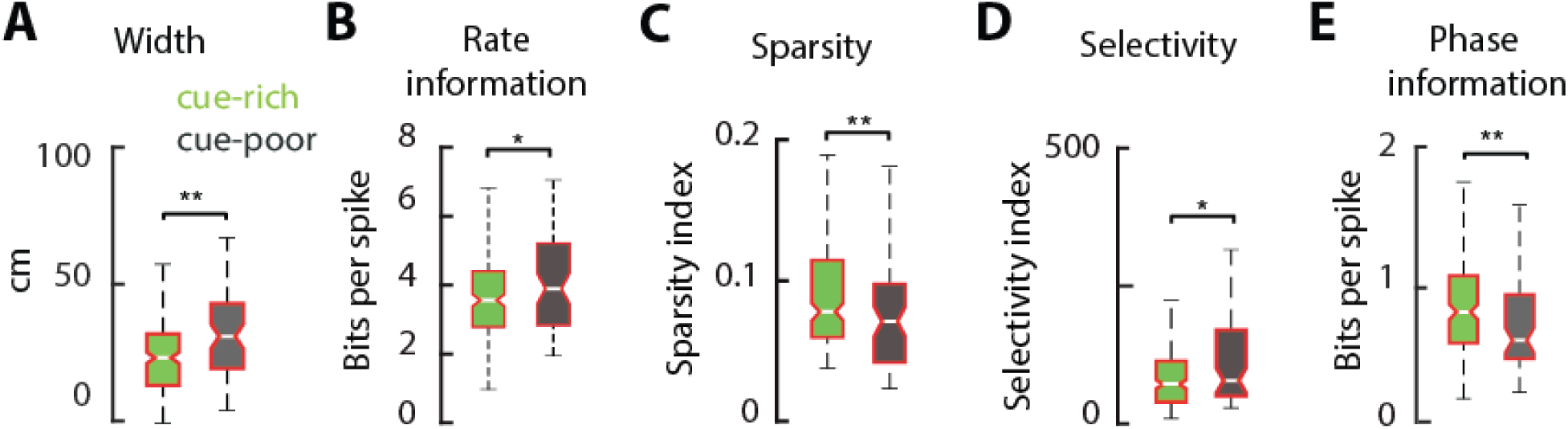
Spatial coding properties of CA1 pyramidal neurons in cue-rich and cue-poor zone after changing reward location. Related to Figures 2 and 3. Place cell metrics in a version of the task in which the location of reward delivery was changed show similar differences as in the main experiment (Figure 2). **A**) Place fields were wider in the cue-poor than in the cue-rich zone (p = 0.0018, Kruskal-Wallis test). **B)** Cue-poor zone place cells carried more spatial information in their firing rates than cue-rich zone place cells (p = 0.021). **C)** Cue-rich zone place cells had sparser (p = 0.0011) and less spatially selective (**D)** firing maps than cue-poor zone place cells (p = 0.020). **E)** Cue-rich zone place cells carried more spatial information in their spike theta-phases than cue-poor zone place cells (p= 0.0036). */**/*** P < 0.05/ 0.01/ 0.001, Kruskal-Wallis test.

**Figure S4.**
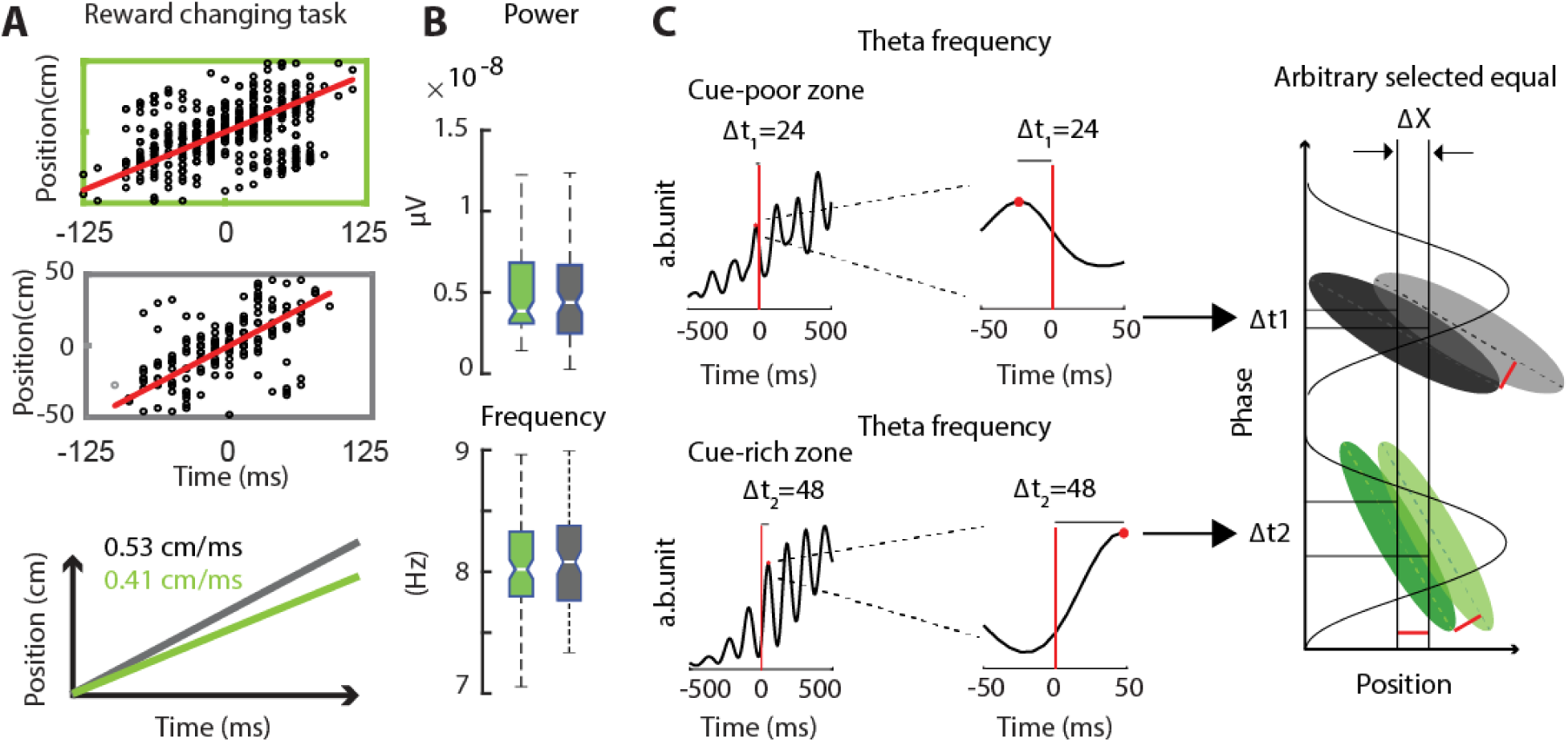
Temporal compression of space in the cue-rich and cue-poor zones. Related to Figure 4. **A**) Correlation between place field peak distance and theta time lag of their spikes for all overlapping CA1 place fields in the cue-rich (left; n = 496) and cue-poor (right; n = 190) zones in version of the task in which reward location was changed. Red line indicates correlation slope (Δs/Δt). Note that compression slope was higher in the cue-poor than in the cue-rich zone (bottom panel), the same trend that was observed in the main experiment (Figure 4). **B**) Theta power and frequency in cue-rich and cue-poor zones were not significantly different. **C**) Schema illustrating how a reduced phase precession slope can explain the increase of temporal compression in the cue-poor zone (grey) compared to the cue-rich zone (green). Plots on the left are theta time scale cross-correlograms of adjacent place field pairs in the cue-poor (top) and cue-rich (bottom) zones. Colored ovals on the right illustrate phase precession spike clouds of the two adjacent place fields in the cue-poor zone (top) and cue-rich zone (bottom).

**Figure S5.**
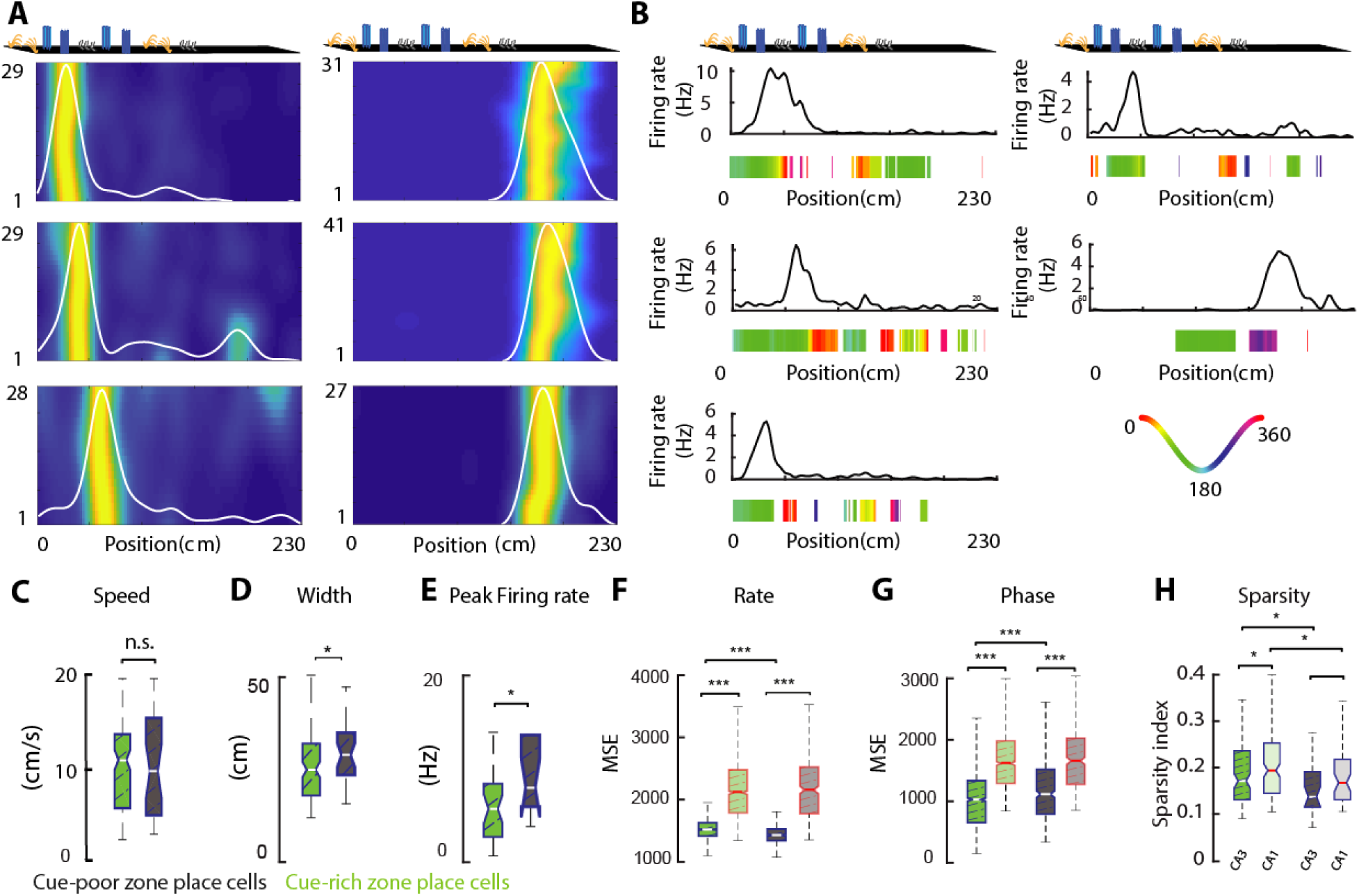
Properties of CA3 place cells in the linear track. Related to Figure 5. **A)** Examples of CA3 place cells recorded on the treadmill. White line is averaged normalized firing rate for the session and color-coded map trial-by-trial firing rate. **B**) Examples of firing rate curves (black lines) and phase maps (color bars) for cue-rich zone CA3 place cells and cue-poor zone place cells. **C**) Average running speed in the cue-rich and cue-poor zones across all sessions with CA3 recordings showed no significant differences (p > 0.05, Kruskal-Wallis test). **D**) CA3 place fields were wider in the cue-poor zone than in the cue-rich zone (p = 0.0136). **E**) Peak firing rate of CA3 place fields were higher in the cue-poor zone than in the cue-rich zone (p = 0.0195). F and G) Comparison of rate and phase decoder performance with chance. */**/*** P < 0.05/ 0.01/ 0.001, Kruskal-Wallis test. **H**) CA3 place cells have less sparse firing rates than CA1 place cells. Clear color boxplots represent CA1 place cells in cue-rich and cue-poor zones.

**Figure S6:**
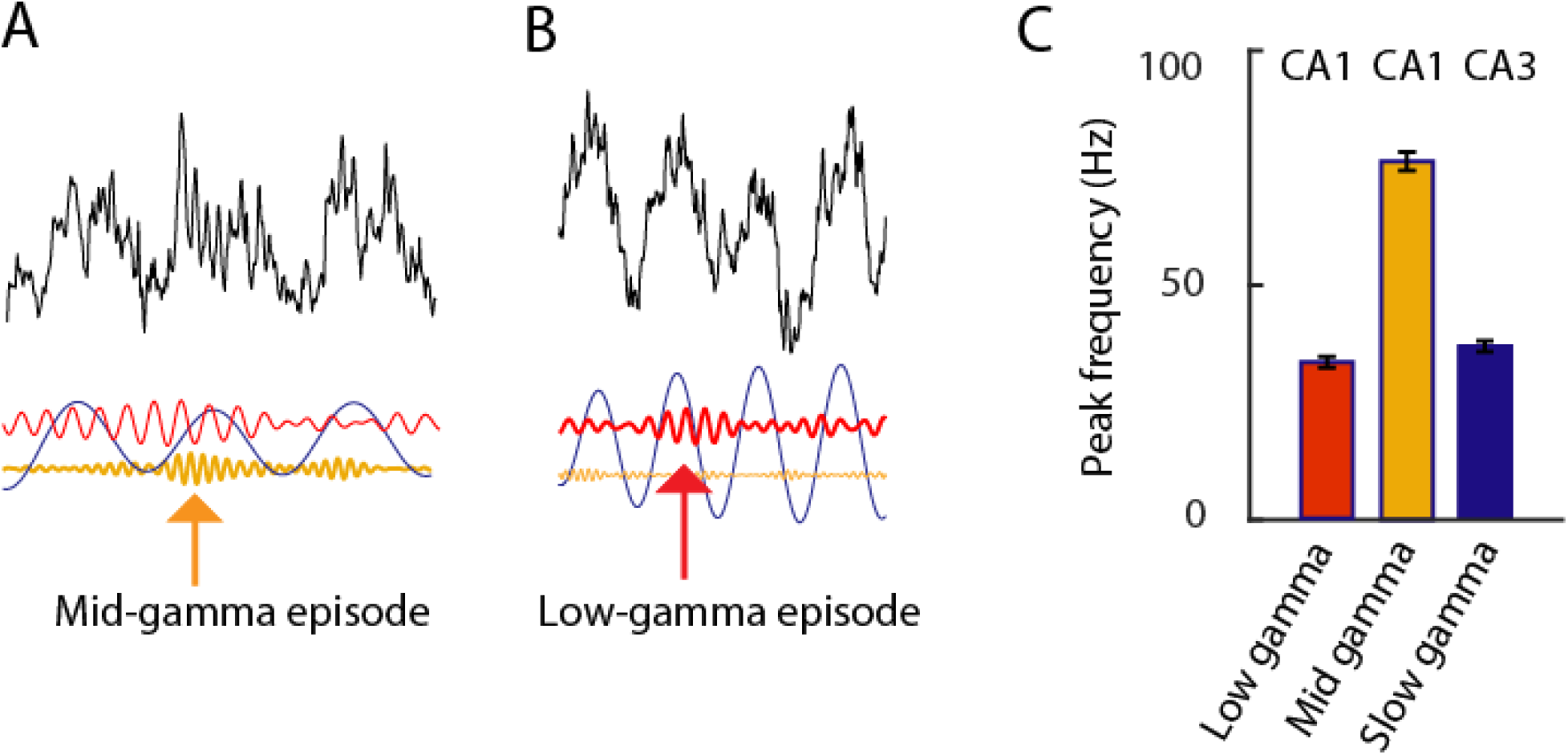
Low and mid-frequency gamma inputs to CA1. Related to Figure 6. **A-B)** Example of wide-band LFP trace (black) and the same segment but filtered in the theta band (5-15 Hz; blue), the slow-gamma (30-60 Hz; red), and the mid-gamma band (60-120 Hz; yellow). Arrow in **A** indicates a mid-gamma episode at the peak of the theta cycle, while arrow in **B** denotes a slow-gamma episode at the descending theta phase. **C**) Peak frequency for all CA1 slow gamma episodes (n = 6 mice), CA1 mid-gamma and CA3 slow gamma.

**Figure S7.**
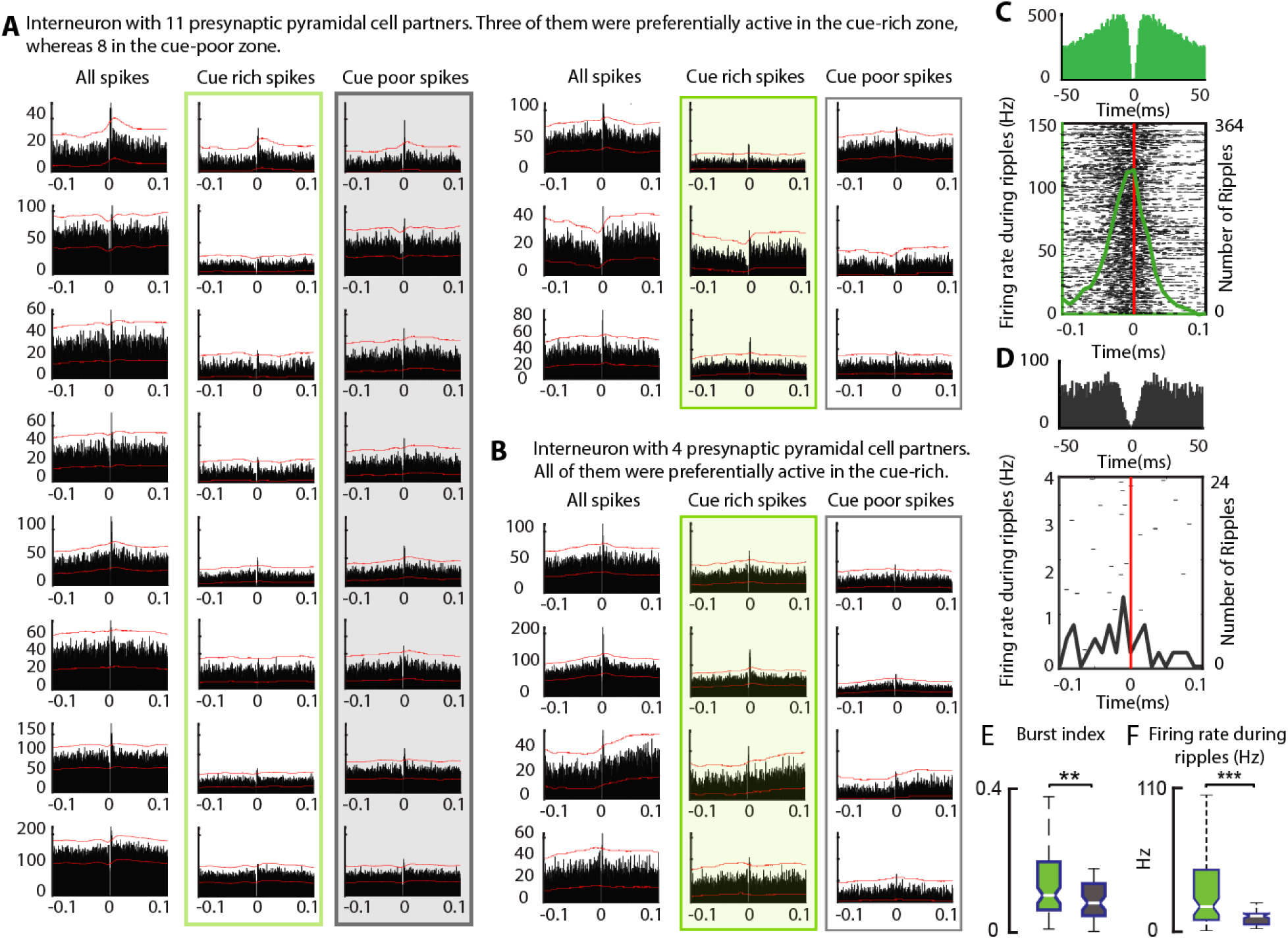
Interneuron recruitment by CA1 pyramidal cells during treadmill running. Related to Figure 7. A) Example of one CA1 interneuron connected to 11 presynaptic CA1 pyramidal cells. First column shows cross-correlograms using all spikes in the recording session, second column using only spikes from the cue-rich zone and third column spikes from the cue-poor zone. Note that in this example, spike-transmission probability was higher in the cue-poor zone than in the cue-rich zone for 8 of the 11 pairs. **B**) Another example of an interneuron connected to 4 presynaptic pyramidal cells. In this case, spike-transmission probability was higher in the cue-rich zone than in the cue-poor zone for the four pairs. **C**) An example of auto-correlogram (top) and peri-sharp-wave ripple firing raster (solid line is average firing across all events) for one interneuron that was preferentially recruited in the cue-rich zone. **D**) Auto-correlogram and peri-sharp-wave ripple firing raster plot for another interneuron that was preferentially recruited in the cue-poor zone. **E**) Interneurons that were preferentially recruited by CA1 place cells in the cue-rich zone (green) were more bursty (p < 0.01, Kruskal-Wallis test) and fired more during sharp-wave ripples (**F**) that interneurons preferentially recruited in the cue-poor zone (grey; p < 0.001).

**Figure S8.**
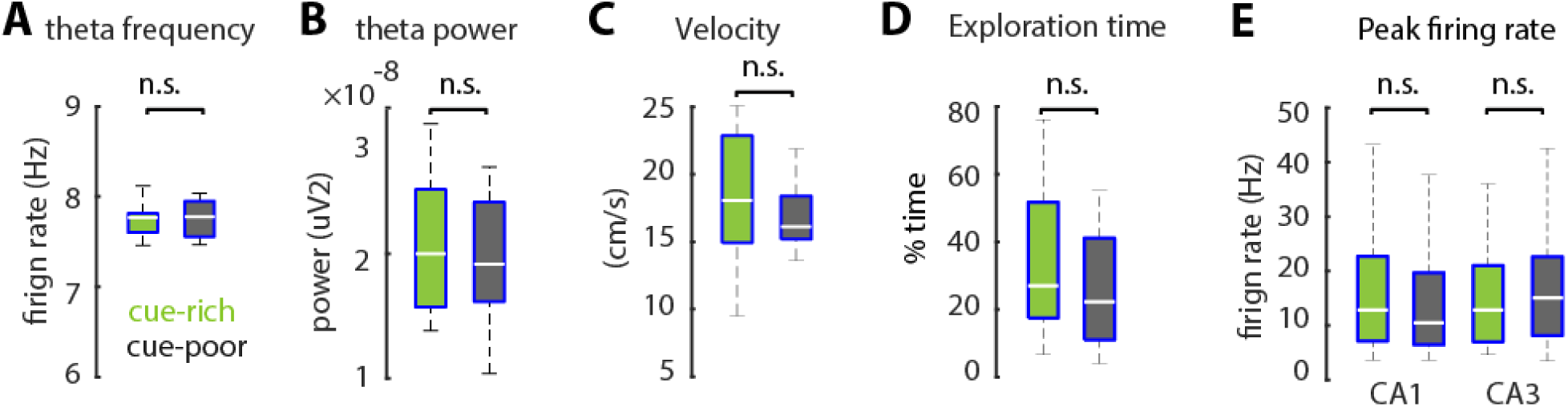
Properties of CA1 and CA3 place cells in the 2D open field. Related to Figure 8. **A)** Mean theta frequency and power (**B**) were not significantly different in the open and object arenas (p > 0.05, Kruskal-Wallis test). **C**) Rats run at similar average speeds in both arenas (p > 0.05). **D**) The proportion of time animals spent actively exploring (velocity > 5cm/s) in the objects and empty arena was not significantly different (p > 0.05). **E**) Peak firing rates of CA1 and CA3 place cells did not differ between the objects and open arenas (p > 0.05).

